# Dengue virus-specific memory B cell subsets differ as a function of infection history

**DOI:** 10.64898/2026.01.30.702935

**Authors:** Tulika Singh, Amir Balakhmet, Jenny M. Granera, Rohan Shinkre, José Victor Zambrana, Nharae E. Lee, Aaron L. Graber, Elias M. Duarte, Sandra Bos, Federico Narvaez, Angel Balmaseda, Eva Harris

**Affiliations:** Division of Infectious Diseases and Vaccinology, School of Public Health, University of California, Berkeley, Berkeley, CA; Division of Immunology and Molecular Medicine, Department of Molecular and Cell Biology, University of California, Berkeley, Berkeley, CA; Sustainable Sciences Institute, Managua, Nicaragua; Laboratorio Nacional de Virología, Centro Nacional de Diagnóstico y Referencia, Ministerio de Salud, Managua, Nicaragua

**Author notes:** Corresponding author: Eva Harris.

## Abstract

The four dengue virus serotypes (DENV1-4) are a major global health threat. Infection generates protective immunity over multiple exposures with different serotypes. Virus-specific memory B cells (MBCs) can contribute to lasting protection, yet their development over multiple DENV infections remains undefined. We comprehensively evaluated frequencies of nine DENV-specific B cell subsets in 58 samples from dengue cases, comparing groups with primary (1°) versus secondary (2°) DENV infection history. Longitudinal sampling from acute infection to 18 months post-symptom onset enabled assessment of DENV-specific MBC temporal dynamics. Critically, we found that DENV-specific B cell frequency differed substantially at the level of phenotypic subsets in 1° versus 2° immunity, despite no significant difference in total frequency of DENV-specific B cells. In particular, DENV-specific IgG+, IgM+, atypical, and class-switched IgD-MBCs were durable until 18 months and accumulated with multiple exposures, representing a *bona fide* memory compartment against DENV. Also, naïve-like IgD+/IgM+ DENV-specific B cells were found. Interestingly, the peak of certain DENV-specific MBCs occurred >3-months post-symptom onset in 2° DENV immunity, suggesting potential for long-term MBC maturation. We demonstrate that DENV-specific MBC subsets differ as a function of infection history, suggesting that 2° DENV immunity does not simply generate a quantitative boost, but a qualitative reprogramming of the memory pool.

**Significance Statement:** The four dengue virus serotypes (DENV1-4) cause the most prevalent human mosquito-borne viral disease. Dengue is a febrile disease that often results in debilitating body pain and can rapidly progress to severe disease involving shock. People are generally protected after multiple exposures to different serotypes. Memory B cells (MBCs) can contribute to lasting protection against subsequent dengue. To understand how these rare DENV-specific MBCs develop over multiple exposures, we compared samples from cases with primary versus secondary (i.e., multiple) DENV infections. We found that instead of a higher frequency of total DENV-specific MBCs, particular subsets of DENV-specific MBCs were higher and peaked later after multiple exposures. This suggests that a qualitative shift in DENV-specific MBCs may contribute to protective immunity.

## Introduction

The four serotypes of mosquito-borne dengue viruses (DENV1-4) co-circulate and cause an estimated 100 million human infections worldwide every year.^1^ Classical dengue is characterized by fever and debilitating body pain, but a fraction rapidly progress to severe dengue disease involving vascular leak.^2^ In endemic regions, people are generally exposed to DENV multiple times, and protective immunity is known to develop after multiple exposures.^3–6^ A key challenge is that a primary (1°) DENV infection generates antibodies that can worsen subsequent disease with a different serotype, whereas secondary (2°) DENV infection is thought to elicit broadly protective immunity across DENV serotypes.^5–11^ Thus, understanding the development of protective immunity over multiple exposures is critical for designing safe and effective vaccines.

Memory B cells (MBCs) generated in response to infection are important because they provide lasting immunity over years and re-activate during subsequent infection to produce a faster response than the initial infection.^12^ A 1° immune response is initiated with naïve B cells that recognize the antigen, whereas a 2° response has the potential to launch from rapidly responding MBCs.^13^ Maturation from naïve to memory generally involves irreversible B cell receptor (BCR) isotype class-switching from IgD to IgM, IgG, and IgA.^14^ DENV-specific MBCs from prior infection are known to contribute to 2° dengue antibody responses.^15–20^ Virus-specific MBCs have been found to harbor a more diverse and cross-reactive repertoire than plasmablasts, offering the possibility of protection against different serotypes of DENV.^16,19,21–23^ Yet, the phenotypes of DENV-specific MBCs poised to respond after 1° versus 2° dengue are unknown.

Recent studies reveal that MBCs with distinct cell surface markers differ substantially in their frequencies, antigen-specific affinity maturation, ability to sense signals of infection, responsiveness to subsequent differentiation, and cell fate upon differentiation.^24–28^ For example, atypical MBCs may be hyporesponsive upon subsequent stimulation and IgM+ MBCs are more likely to engage in germinal center reactions, whereas IgG+ MBCs have a higher propensity to differentiate into plasmablasts (PBs).^29,30^ Thus, it is possible that distinct phenotypes of DENV-specific MBCs are present after 1° versus 2° dengue that confer distinct downstream functions to modulate infection. Yet, the immunophenotyping of DENV-specific MBC subsets has been lagging due to the enormous technical difficulty of enumerating rare DENV-specific B cells.^31^ Most of these studies have focused on measuring total DENV-specific B cells (not subsets) or DENV-specific MBCs of a single type.^17,20,31–40^ Thus, there is a gap in knowledge regarding the development and durability of distinct DENV-specific MBC subsets after 1° versus 2° dengue.

Here, we compared 1° versus 2° DENV B cell responses by developing a modified approach to detect low frequencies of DENV-specific B cells, immunophenotyped DENV-specific MBC subsets from 58 pediatric samples and evaluated longitudinal DENV MBC dynamics up to 18 months post-infection. Collectively, the data reveal a qualitative reprogramming of the DENV-specific MBC compartment as a function of infection history.

## Results

### Development of a modified approach for detection of lasting DENV-specific B cells

We modified pre-existing approaches to detect flavivirus-reactive B cells by flow cytometry with the goal of improving detection of lasting DENV-specific B cells.^20,32,41^ First, we grew viruses on Vero-furin cells to enrich for mature viral particles, since potent neutralizing antibodies are known to target quaternary epitopes and mature virions.^20,32,40,42–44^ Second, we used viral strains derived from our Nicaraguan study population as probes and included multiple frequently circulating serotypes (DENV1, 2, and 3) to maximize detection of DENV-specific B cells.^45^ Third, each virus was singly labelled with Alexa Fluor (AF)-488 and AF647 and then assembled into a six-antigen cocktail for staining PBMCs (Fig 1A). DENV-specific cells were thus defined as the subset of live B cells that react with both DENV-AF488 and DENV-AF647 (i.e., double-positive), as prior experience indicates that single-positive B cells may be only 50% virus-reactive (Fig 1B).^41^ Labelled antigens ranged between 10^5^-10^7^ viral particles per mL and demonstrated binding to monoclonal antibodies (mAbs) against known quaternary epitopes (Fig S1, S2). Titration of the pooled antigens on a DENV-immune versus a DENV-naïve PBMC sample, yielded an optimal staining volume of 7.5uL antigen per million PBMCs (Fig S3). This condition rendered a frequency of 20 DENV-specific cells per 10,000 B cells in the immune sample, which was higher than the maximal non-specific background of 5 DENV-specific cells per 10,000 B cells in the no-antigen control and DENV-naïve sample (Fig S3). Thus, this pooled antigen condition was used for the rest of this study.

**Figure 1.**
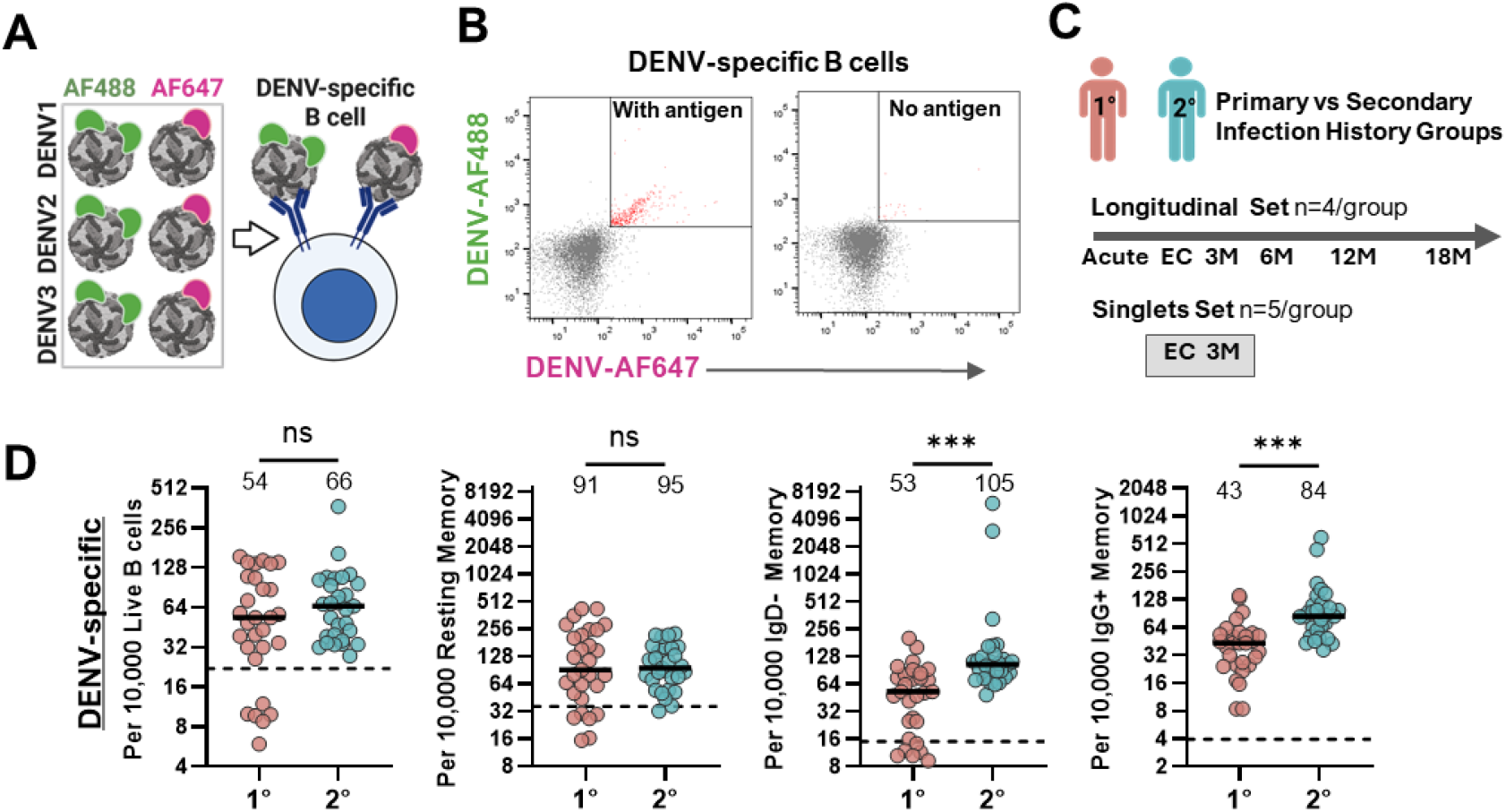
Enumerating DENV-specific B cells from longitudinal samples of primary versus secondary dengue cases reveals different frequencies by subsets and not total. **A**. Our modified strategy for defining DENV-specific B cells involved labelling DENV1, DENV2, and DENV3 viruses each with Alexa Fluor (AF)-488 and AF647 and combining these six labelled viruses into an antigen cocktail for staining PBMCs, such that DENV-specific B cells were defined as double-positive for both AF488 and AF647 DENV antigen. **B**. Total DENV-specific B cells were defined as CD3-/CD14-/CD16-/Viable/CD19+/AF488+/AF647+ double positive (red dots). Shown are double-positive cells in a DENV-immune Nicaraguan pediatric PBMC sample with antigen versus a Nicaraguan adult PBMC sample without antigen, which served as the lower limit of detection. **C**. Study design for comparison of 1° versus 2° DENV B cell responses in pediatric dengue cases included a set of longitudinal PBMC samples and another single-timepoint set for a total of 58 samples. In the Longitudinal Set, participants with 1° (n=4) and 2° (n=4) DENV infection histories were sampled over time at acute, EC, 3M, 6M, 12M, and 18M. The Singlets Set included one sample per 1° (n=5) and 2° (n=5) dengue participant collected either at EC or 3M. **D**. Frequency of DENV-specific B cells in all 1° (pink) versus 2° (blue) dengue samples irrespective of timepoint. DENV-specific cells per 10,000 total live B cells, resting MBCs, class-switched IgD-MBCs, and IgG+ MBCs. Median shown with solid line and with numeric value per group. Dotted line is limit of detection based on mean of adult PBMC samples without antigen (n=7). One-tailed Mann-Whitney test, p<0.001 (***).

### Participant characteristics

We analyzed a total of 58 peripheral blood mononuclear cell (PBMC) samples from 18 pediatric patients with DENV infection (Table S1 and S2). These participants were selected based on sample availability as a convenience set from our hospital-based observational study in Managua, Nicaragua. We compared the DENV-specific B cell profile of primary (1°; n=9) versus secondary (2°; n=9) dengue cases. A 1°DENV infection history was classified by a DENV antibody inhibition ELISA (iELISA) titer below 2560 in convalescent-phase samples, and 2°infections were classified by titers of 2560 or higher in convalescent-phase samples, as before.^46,47^ From 8 patients, we immunophenotyped longitudinal PBMCs collected during acute infection, 2-4 weeks post-symptom onset (early convalescence; EC), as well as 3, 6, 12, and 18 months (M) post-symptom onset (i.e., Longitudinal Set; Fig 1C and Table S1). From 10 patients, we immunophenotyped a single timepoint at EC or 3M (i.e., Singlets Set; Fig 1C and Table S2), which we reasoned may be the peak DENV-specific B cell response.

### Frequency of DENV-specific B cells

On average, we analyzed ∼1,000,000 PBMCs, ∼100,000 live B cells (CD3-/CD14-/CD16-/CD19+), and 561 DENV-specific cells per sample, with no significant differences in raw counts by infection history (n=58; Fig S4). Several B cell subsets were defined using additional conventional markers: naïve B cells (CD20+/IgD+), IgD+/IgM+ naïve B cells (CD20+/IgD+/IgM+), class-switched MBC (CD20+/IgD-), activated MBC (CD20+/IgD-/CD27+/CD21-), resting MBC (CD20+/IgD-/CD27+/CD21+), atypical MBC (CD20+/IgD-/CD27-/CD21-), IgG+ MBC (CD20+/IgD-/IgG+), IgM+ MBC (CD20+/IgD-/IgM+), and IgD-/IgM-/IgG-MBC (Fig S5, Table S3).^27^ Within total live B cells and each of the nine B cell subsets, DENV-specific cells were enumerated as AF488+/AF647+ double-positive. Frequency of DENV-specific B cells was calculated per 10,000 parental B cells. Pooling all timepoints within each 1° and 2° group, there were no significant differences by infection history in frequencies of total DENV-specific B cells (median: 1°=54, 2°=66 per 10,000 live B cells) or resting MBCs (median: 1°=91, 2°=95 per 10,000 resting MBCs; Fig 1D). DENV-specific class-switched IgD-MBCs (median: 1°=53, 2°=105 per 10,000 IgD-MBC) and IgG+ B cells (median: 1°=43, 2°=84 per 10,000 IgG+ B cells) were significantly higher upon 2° than 1° infection, as expected (p<0.001; Fig 1D). Thus, enumeration of DENV-specific B cell subsets provides distinct information from the total frequency and distinguishes 1° versus 2° dengue immunity.

### Differences in early and lasting DENV MBC subsets by infection history

To understand the generation and durability of DENV-specific B cell responses in 1° versus 2° dengue, we compared frequencies at acute, EC, and 18M timepoints, as these represent immunologically relevant phases of initiation, establishment of memory, and retention of memory (Fig 2). At all timepoints, there were no significant differences in the frequency of total DENV-specific B cells by infection history (Fig 2A, B, S6). In acute infection, DENV-specific IgG+ MBCs were not significantly different, but DENV-specific IgM+ MBCs were significantly higher in 2° than 1° dengue (p<0.05, n=4 per group; Fig 2C, D). Transient circulation of plasmablasts (PBs; CD38+/CD27+) during infection constitutes an immune response. There were no significant differences in total PBs by infection history (Fig 2E, F). We then quantified IgM+ versus IgM-PBs since IgM+ B cell receptors (BCR) are maintained on PB cell surfaces but IgG+ BCR expression is variable (Fig 2G).^48^ We found that the 2° dengue group possessed significantly higher IgM-PBs compared to the 1° group (p<0.05, Fig 2H). Thus, higher frequencies of DENV-specific IgM+ MBCs and IgM-PBs distinguished acute 2° from 1° dengue B cell responses.

**Figure 2.**
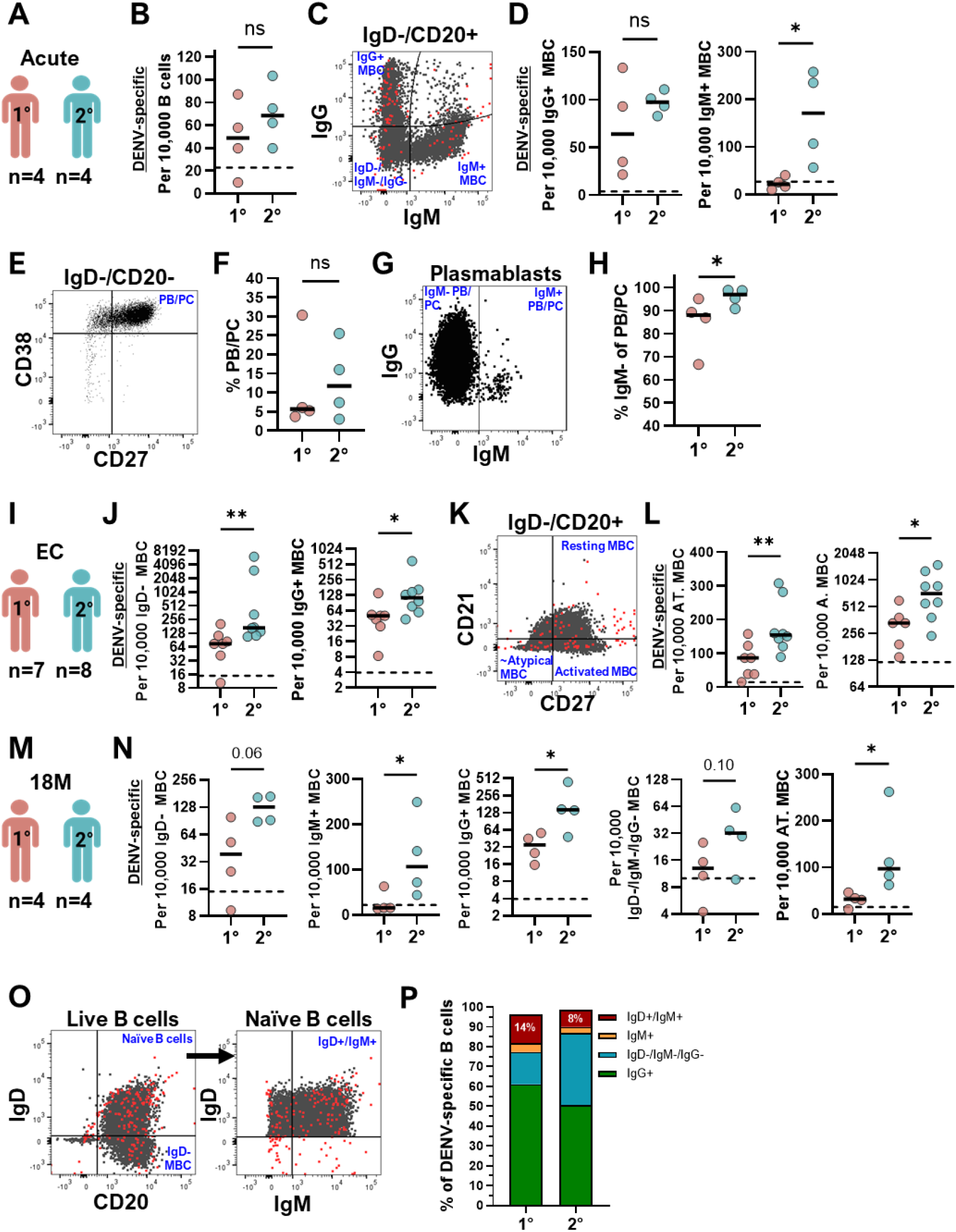
Differences in early and lasting DENV-specific B cell responses in primary versus secondary dengue cases. **A**. Comparisons of <u>acute</u> 1° (n=4; pink) versus 2° (n=4; blue) B cells. **B**. Total DENV-specific B cell frequencies. **C**. From total class-switched MBCs (IgD-/CD20+), the IgG+, IgM+ and IgM-/IgG-subsets were defined, as well as the DENV-specific B cells (red) within each subset. The IgD-/IgM-/IgG-subsets may largely represent IgA+ B cells. **D**. Frequency of DENV-specific cells per 10,000 IgG+ and IgM+ MBCs in 1° versus 2° dengue cases. **E**. PB/PCs (plasmablasts including plasma cells) were gated as CD38+/CD27+ from the IgD-/CD20-parental population. **F**. PB/PCs as a percentage of total B cells in the 1° versus 2° acute DENV infection. **G**. Gating of PB/PCs subsets as IgM+ versus IgM-, which represent largely IgG-expressing PB/PCs with variable cell surface BCR. **H**. IgM-PB/PCs as a percentage of total PB/PCs in 1° versus 2° acute DENV infection. **I**. Comparisons of 1° (n=7) versus 2° (n=8) dengue cases at <u>early convalescence</u>. **J**. Frequency of DENV-specific B cells per 10,000 parental populations of class-switched IgD-/CD20+ and IgG+ MBCs in 1° versus 2° dengue cases. **K**. From total class-switched MBCs (IgD-/CD20+), resting MBCs (CD21+/CD27+), activated MBCs (CD21-/CD27+), and a double-negative subset that includes atypical MBCs (CD21-/CD27-) were defined, from which DENV-specific B cells (red) were quantified. **L**. Frequency of DENV-specific cells per 10,000 parental atypical MBCs (AT. MBCs) or per 10,000 parental activated MBCs (A. MBC) subsets in 1° versus 2° dengue cases. **M**. Comparisons of <u>18M</u> DENV-specific B cells from 1° (n=4) versus 2° (n=4) dengue cases. **N**. Frequency of DENV-specific B cells per 10,000 parental populations of class-switched IgD-/CD20+, IgM+, IgG+, IgD-/IgM-/IgG-, and atypical (AT.) MBCs in 1° versus 2° dengue cases. **B-N**. For all dot plots, solid line at median is shown, alongside One-sided Mann Whitney tests with significant differences indicated as p<0.05 (*) and p<0.01 (**), trending values (p≤0.1) numerically indicated, and non-significant differences labeled as ns. Dotted line indicates limit of detection based on mean of adult PBMC samples without antigen for each subset (n=7). **O**. From total live B cells, naïve B cells (IgD+/CD20+) were gated; within this subset, the vast majority of cells were also IgM+ (representative gate shown from 1° dengue case 18-month sample). DENV-specific B cells in red. **P**. Distribution of BCR isotypes among DENV-specific B cells at 18M. Mean proportion of IgG+ (green), IgM+ (orange), IgD+/IgM+ (red), and IgD-/IgM-/IgG-(blue) B cells as a proportion of DENV-specific B cells in 1° (n=4) versus 2° (n=4) dengue cases.

Among EC samples (1° n=7, 2° n=8), 2° dengue cases demonstrated significantly higher DENV-specific IgD-class-switched MBCs (p<0.01) and IgG+ B cells (p<0.05) than 1° cases (Fig 2I, J). MBC phenotypes were also assessed via differential CD21 and CD27 expression (Fig 2K). DENV-specific atypical and activated MBC subsets were significantly higher in 2° than 1° dengue groups at EC (p<0.01 and p<0.05, respectively; Fig 2L). These features are commensurate with greater responsiveness and/or formation of MBCs in 2° than 1° dengue immunity.

The persistence and accumulation of virus-specific cells with multiple exposures is a classic hallmark of memory cell types. At 18M, many DENV-specific subsets were detectable above the level of background. Specifically, there were significantly higher frequencies of IgG+, IgM+, and atypical MBCs in the 2° than 1° dengue group (p<0.05, Fig 2M, N). Meanwhile, 2° IgD-class-switched and IgD-/IgM-/IgG-MBCs trended towards significance (p=0.06 and p=0.1, respectively; Fig 2N). While IgG+ MBCs are largely studied in dengue immunity, our findings indicate that DENV-specific IgM+ B cells and atypical MBCs may also have a role in long-term dengue immunity.

### Persisting naïve-like DENV-specific B cells

Oddly, a population of naïve-like B cells (CD20+/IgD+) also contained many DENV-specific cells (Fig 2O). The majority of this DENV-specific naïve-like subset was IgD+/IgM+ and above the detection limit, whereas the IgD-only (IgD+/IgM-) subset was largely below the detection limit, thus the IgD-only subset was not analyzed further (Fig S7). To conservatively estimate the portion of IgD+/IgM+ cells among DENV-specific B cells at 18M, we developed another gating strategy by sub-sampling bright double-positive DENV-specific B cells and tailoring the cut-off for each isotype (i.e., IgD, IgM, and IgG) to each individual’s isotype distribution from total B cells (Fig S8). We found that IgD+/IgM+ B cells constitute on average 14% and 8% of all DENV-specific B cells at 18M after 1° and 2° dengue, respectively (Fig 2P).

### Distinct dynamics of DENV-specific subsets by infection history

We then sought to define whether the trajectory of each DENV-specific B cell subset differed by 1° versus 2° infection history (n=4 individuals per group and n=6 timepoints; Fig 3A, S9). Using a Two-way Repeated Measures ANOVA, accounting for individual variability, we tested for global differences in the average DENV-specific B cells by 1) infection history across all0020timepoints; and 2) only timepoint regardless of infection history (Fig 3A). Again, no significant differences were observed for total DENV-specific live B cells (Fig 3A). However, there was a significant effect of 1° versus 2° infection history on DENV-specific subsets, namely class-switched IgD-, activated, IgG+, IgD-/IgM-/IgG-, and atypical MBC subsets, where 2° dengue cases demonstrated a higher response than 1° cases (p<0.01, Fig 3A). Interestingly, atypical MBCs (CD27-/CD21-), but not resting MBCs (CD27+/CD21+), differed by infection history. DENV-specific B cell subsets that additionally demonstrated significant differences by timepoint, and not just infection history, included class-switched IgD-, activated, and atypical MBCs, which are characterized by a high average peak response at EC (p<0.05; Fig 3A). Importantly these differences were not observed in the overall parent B cell compartment frequencies, reinforcing that these distinct dynamics are specifically occurring in the anti-DENV response (Fig S10, S11).

**Figure 3.**
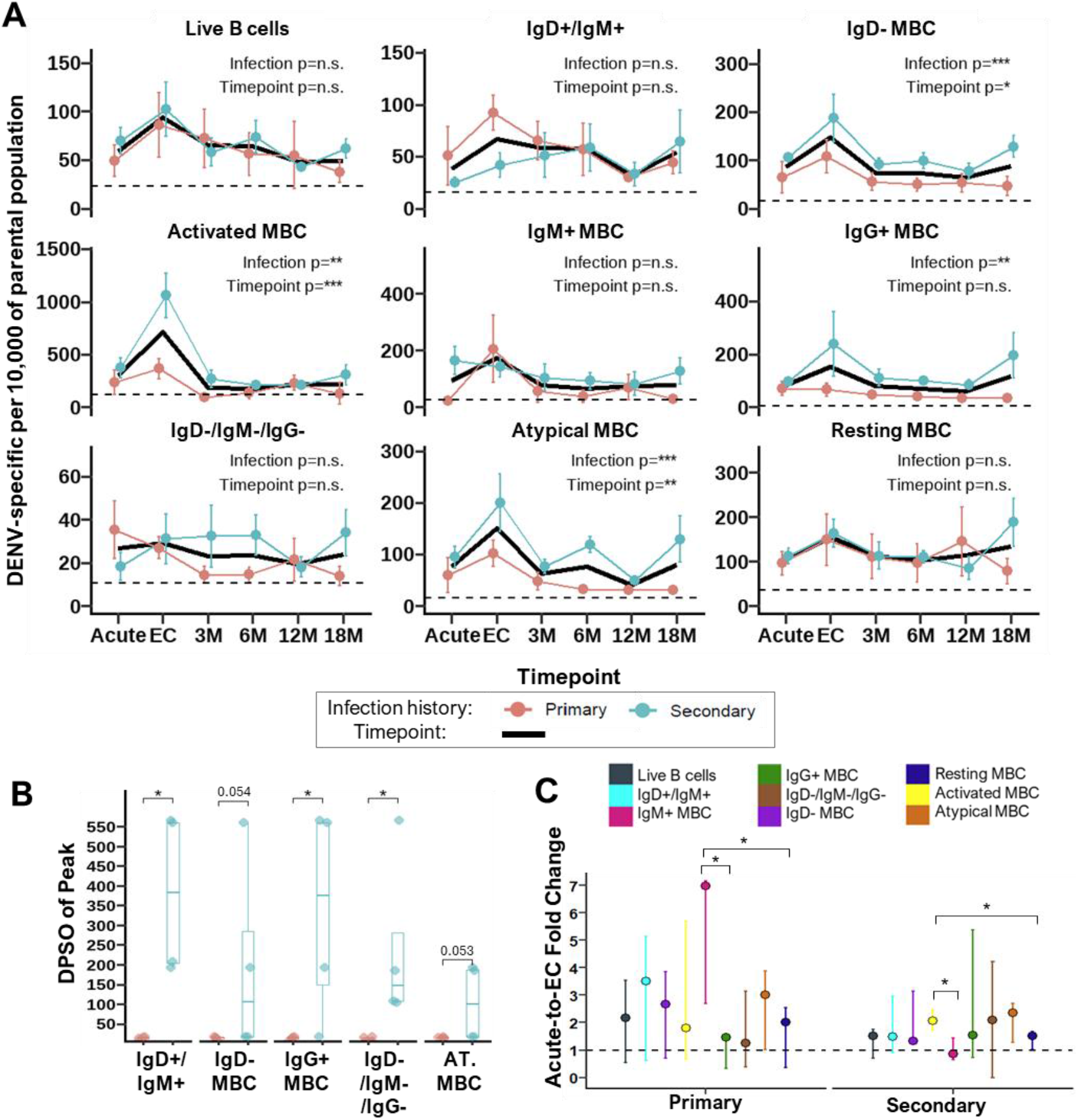
Distinct dynamics and peaks of each memory subset by immune history. **A**. Two-way Repeated Measures ANOVA was conducted to evaluate the main effect of timepoint and infection history on DENV-specific B cell frequency in the 1° (n=4 individuals and n=6 timepoints; pink) versus 2° (n=4 individuals and n=6 timepoints; blue) infection history groups, accounting for individual variability. Timepoint was treated as a within-subject repeated measure, and infection history group (i.e., infection) was treated as a between-subjects measure. Mean frequency of DENV-specific B cells per 10,000 parental cells at each timepoint (Acute, EC, 3M, 6M, 12M, and 18M) is shown with standard error of the mean for each subset. The results of the test are annotated on each panel, showing the main effect significance for Infection and Timepoint (*, p<0.05; **, p<0.01). The solid black line represents the main effect of timepoint, combining 1° and 2° infection groups. Dotted line indicates limit of detection based on mean of samples without antigen for each subset (n=7). **B**. Day post-symptom onset (DPSO) of the peak indicates the day of the peak frequency of DENV-specific B cells per 10,000 parental cells from the six timepoints measured for each participant. Box plot indicates median and range of each group. One-sided Wilcoxon signed rank test used to compare groups of 1° (n=4) and 2° (n=4) dengue cases, where significant differences are indicated by * (p<0.05) and trending p-values with number. **C**. Fold-change in DENV-specific B cells from acute to EC represents the response to infection and was calculated per individual using frequencies per 10,000 parental cells, where a value of 1 indicates no difference (dotted line). Median and range of fold-changes are shown for each 1° (n=4) and 2° (n=4) dengue group by subsets, indicated in color. Pairwise two-sided Wilcoxon signed rank test was applied to assess differences in fold-change by subset with significance as p<0.05 (*).

Unexpectedly, in 2° dengue cases, we observed an uptick from 12M to 18M in DENV-specific IgD-, IgG+, IgD-/IgM-/IgG-, and atypical MBCs, unlike the 1° cases (Fig 3A, S9). This uptick was not present in the overall parent B cell compartment indicating that this is not a general B cell trend but specific to responses against DENV (Fig S11). Since our analysis of DENV-specific cells per 10,000 parental cells reflects a relative metric within each subset (Fig S9), we cross-checked that this 12-18M uptick was also present using an absolute metric of DENV-specific cells as a percentage of live B cells (Fig S12). Indeed, three out of four 2° dengue cases demonstrated this 12-18M uptick (Fig S9, S12), suggesting potential for long-term DENV-specific B cell maturation, or fluctuations from MBCs transiting between tissues and the periphery in 2° immunity.

### A delayed peak in DENV-specific B cell frequency in secondary cases

Next, we sought to evaluate the timing of peak DENV-specific MBC responses in PBMCs. For each 1° and 2° dengue case evaluated longitudinally (n=4 individuals per group), the peak response was defined as the day post-symptom onset of the highest DENV-specific B cell frequency per 10,000 parental cells. Surprisingly, three DENV-specific B cell subsets peaked significantly later in 2° than 1° dengue immune responses (p<0.05), and two were trending towards significance (p<0.06; Fig 3B). This is consistent with longer-term development of DENV-specific MBCs in 2° than 1° immunity.

Given this delayed peak, we sought to quantify the extent of DENV-specific B cell variability that accounts for an immune response. Since a viable response occurred between acute infection to resolution of disease at EC, we quantified the fold-change in DENV-specific B cell frequencies from acute to EC (Fig 3C). In the 1° response, DENV-specific IgM+ MBCs demonstrated the greatest fold-change (7-fold), whereas in the 2° response activated, atypical and IgD-/IgM-/IgG-MBCs demonstrated the greatest fold-change (2.3-, 2.3-, and 2.1-fold respectively; Fig 3C). By infection history, fold-change in 1° IgM+ and 2° activated DENV-specific MBCs was significantly greater than that observed for 1° IgG+, 1° resting, 2° IgM+, and 2° resting MBCs, suggesting differences in the early B cell responsiveness based on phenotypic subset (p<0.05; Fig 3C).

### Inferring cellular processes in primary and secondary DENV immune responses

Distinct cellular maturation processes underlie fundamental differences in 1° versus 2° immune responses. To estimate these processes in DENV-specific B cell responses, we tested for direct (i.e., positive) correlations in DENV-specific B cell subset frequencies between each consecutive timepoint and plotted significant correlations (alpha = 0.1; n=4 individuals per group) as a directed network graph (Fig 4). To enrich plausible cellular processes, correlations of known biologically implausible processes were omitted (Table S4, Fig S13). As expected, the majority of correlations observed in DENV-specific B cell frequencies were not found in total B cell frequencies, reinforcing that the correlations identified were specific to the anti-DENV response (Fig S14).

**Figure 4.**
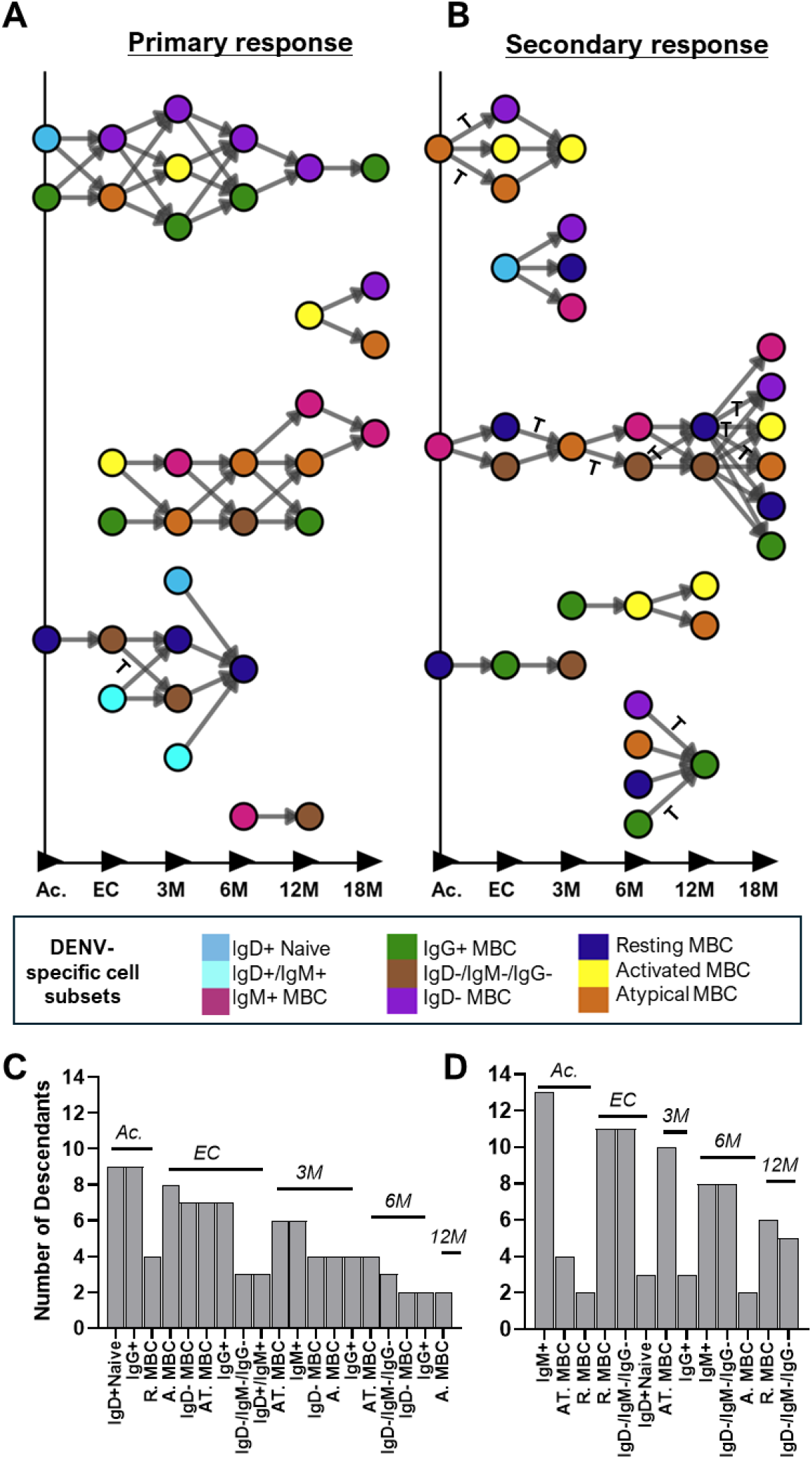
Inferring cellular processes based on direct correlations over consecutive time bins in primary and secondary DENV immunity. Direct Spearman Rank correlations of DENV-specific B cell frequencies per 10,000 parental cells between consecutive timepoints (i.e., Acute-EC, EC-3M, 3M-6M, 6M-12M, and 12M-18M) at p<0.1 were plotted as an arrow on a directed network graph for each (**A**) 1° (n=4 participants; and (**B**) 2° (n=4 participants) dengue groups. Each arrow represents a hypothesized biologically plausible cellular transition or a co-incident and independent process. If known biologically <u>im</u>plausible B cell transitions were correlated at p<0.1, these arrows were excluded to visualize biologically plausible B cell transitions (see Supplement). Colored circles indicate DENV-specific B cell subsets. Correlations also found in the total B cell frequencies were annotated on or adjacent to the arrow with “T.” **C**. From this directed network graph, descendants of each DENV-specific B cell subset were enumerated to evaluate influential cell types in 1° (n=4 participants) and (**D**) 2° (n=4 participants) DENV immune responses. The value atop bar plots indicates the timepoint (Ac is Acute). DENV-specific B cell subsets abbreviated as follows: activated MBCs (A. MBC), resting MBC (R. MBC), atypical MBCs (AT. MBC), IgD-/IgG-/IgM-(inferred IgA), class-switched MBCs (IgD-MBC), IgD-/IgG+ MBCs (IgG+), IgD-/IgM+ MBCs (IgM+), and naïve-like B cells (IgD+ naïve).

First, we examined differences in the initiation of the 1° versus 2° DENV-specific B cell response. Acute DENV-specific IgD+ naïve B cells gave rise to the biggest network in the 1° but not the 2° response, as expected (Fig 4A). Interestingly, in 2° dengue, acute DENV-specific IgM+ MBCs were associated with the largest network (Fig 4B). A positive correlation of 2° acute and 3M DENV-specific atypical MBCs was found with later IgD-class-switched and activated MBC levels, suggesting that atypical MBCs are capable of responding in DENV infection (Fig 4B). We then assessed influential DENV-specific B cell subsets in the 1° versus 2° responses by enumerating the number of descendants in downstream correlations. Acute and early DENV-specific IgD+ naïve and IgG+ B cells were most influential to 1° immunity, whereas IgM+, atypical, resting, and IgD-/IgM-/IgG-MBCs were most influential in 2° dengue immunity (Fig 4C, D).

Surprisingly, some significant correlations in DENV-specific B cell subset frequencies were found between 12M and 18M, indicating the presence of non-random long-term processes in the development of DENV-specific MBCs (Fig 4A, B). Most of these correlations did not overlap with long-term correlations in total B cell frequencies (Fig 4, S14). In both 1° and 2° groups, these 12-18M correlations emerged from IgD-class-switched B cell subsets (Fig 4A, B). Also, the frequency of IgM+ and resting MBCs was correlated with a later high frequency of the same subsets, indicating potential for self-replenishment (Fig 4A, B). Overall, these correlations reveal a hypothetical working model for the long-term development of human DENV-specific MBCs upon 1° versus 2° dengue.

## Discussion

MBCs are a key component of long-lived immunity, yet their heterogeneity and development remain poorly understood due to the technical challenge of measuring these rare cells in limited human samples. This challenge is particularly pronounced in the context of DENV, where interlaced viral envelope proteins present complex epitopes that may not be adequately recapitulated with recombinant protein antigens for DENV-specific B cell detection. In DENV infections, protective immunity is generated after exposure to different serotypes in endemic settings. To understand the development of DENV-specific MBCs over multiple exposures, we phenotyped 58 samples from pediatric dengue cases and compared 1° versus 2° immunity longitudinally up to 18 months post-infection. This in-depth evaluation offers several conceptual insights into the establishment and development of DENV-specific MBCs in human immunity. First, we show that 2° DENV immunity is driven by a *qualitative* rather than *quantitative* shift in the MBC compartment, as differences in frequency from 1° immunity were observed exclusively among phenotypic subsets (i.e., atypical, IgG+, IgM+, class-switched IgD-MBCs) but not in total DENV-specific B cell frequency. Second, we found that these subsets were durable and accumulated with multiple exposures, representing a *bona fide* memory compartment against DENV. Notably, we identify DENV-specific atypical and IgM+ MBCs, which represent understudied subsets of memory. Third, longitudinal analysis revealed that atypical, IgG+, and class-switched IgD-MBC subsets differ in dynamics by infection history, reinforcing differences in MBC selection by exposure. Surprisingly, we observed delayed peaks of several DENV-specific B cell subsets in 2° but not 1° responses, suggesting potential for long-term MBC maturation in 2° immunity. Lastly, we then built a hypothetical working model of DENV-specific MBC development using temporal correlations, which posits and IgD+ naïve B cells and IgG+ MBCs as early responders in 1° immunity and atypical and IgM+ MBCs as early responders in 2° immunity. Altogether, this suggests that development towards protective immunity with multiple DENV exposures may be driven by selection of qualitatively different MBC subsets that are poised to respond in subsequent exposure.

Estimating the frequency of DENV-specific MBCs in circulation has been an important goal of many studies, as higher frequencies of MBCs are thought to lead to more robust subsequent immune protection. We find that DENV-specific cells constitute 0.72% of all B cells and 0.78% of class-switched IgD-MBCs, on average, among pediatric dengue patients. Our frequencies are concordant with prior studies reporting 0.3-0.9% DENV-specific B cells using flow cytometry approaches.^20,32,34^ Prior estimates of DENV-specific B cell frequency using B cell stimulation or immortalization varied widely from 0.01-16% of B cells.^17,38,39,49–51^ Differences in estimates by technique may be due to low immortalization efficiency of B cells *in vitro*, some proliferation *in vitro*, and varied optimal culture conditions for different B cell subsets.^52–56^

Since 1° DENV infection history is associated with subsequent risk of immunopathology, and 2° DENV infection histories are thought to achieve broad protection,^5,6,10,11^ we compared the distinguishing features of 1° versus 2° DENV MBCs. While there was no significant difference in the frequency of total DENV-specific B cells, IgG+ and class-switched IgD-MBC subsets were higher in 2° than 1° dengue immunity, supporting that 2° immunity is characterized by the accumulation of these subsets. Class-switched IgD-and IgG+ MBCs can differentiate into PBs or re-engage in germinal center reactions for affinity maturation.^13^ Accordingly, we found significantly higher IgM-PBs (likely IgG+) in acute 2° than 1° dengue, similar to prior reports.^57–59^ Also, the peak of DENV-specific activated MBCs observed in 2° but not 1° EC samples reinforces that pre-existing MBCs contribute to subsequent DENV responses, supporting prior work.^18,37,38^

Yet, in addition to sequential re-engagement of IgG+ MBCs, our hypothetical working model indicates potential for replacement and diversification of the DENV-specific IgG+ B cell compartment from other DENV-specific memory compartments during 2° immunity – such as from IgM+, resting, and atypical MBCs. This is supported by a prior study indicating limited further affinity maturation in flavivirus immunity, but rather diversification through pre-existing cross-reactive clones.^15,60^ Such replacement of clones is compatible with the modest levels of somatic mutations found in several flavivirus neutralizing monoclonal antibodies.^61^

Interestingly, we found that only DENV-specific IgM+ B cells were significantly higher in acute 2° than 1° dengue, suggesting that IgM+ B cells may be recalled from memory. While the existence of IgM+ MBCs is debated, several studies have found the existence of virus-specific IgM+ B cells against Zika virus, yellow fever vaccine, SARS-CoV-2, and rotavirus, underscoring that this is a *bona fide* memory compartment.^22,41,62,63^ Intriguingly, a prior study even found that DENV-specific IgM+ B cell clones were expanded and mutated in dengue patients, reinforcing their potential to respond to DENV infection.^20^ IgM+ B cells tend to differentiate to germinal center cell fates *in vitro* whereas IgG+ B cells demonstrate greater propensity to differentiate into PBs, suggesting that the DENV-specific IgM+ MBC compartment may confer distinct functions than IgG+ MBCs.^30^ Importantly, the differences in magnitude observed in the DENV-specific IgD-/IgM+ subset were not present in the naïve-like DENV-specific IgD+/IgM+ subset, underscoring that these are two distinct populations.^64^ Thus, IgM+ MBCs represent an underappreciated compartment that contributes to rapid responses in 2° DENV infections.

Interestingly, naïve-like virus-specific B cells (IgD+/IgM+) have been sporadically identified across a few pathogens, and our work provides evidence of their existence in the context of DENV infections. Virus-specific naïve-like IgD+/IgM+ B cells were also found to be long-lasting in the vaccine response against the related yellow fever flavivirus.^62^ Prior studies have found virus-binding naïve-like B cells in the context of SARS-CoV-2 and influenza virus infections, which similarly peaked at EC.^65–69^ Some studies have found that naïve-like B cells contribute to transient IgG1 PBs, autoreactive specificities, weakly neutralizing specificities, severe disease, and anergic cell phenotypes.^65–67,70^ Others found that such naïve-like B cells provided a crucial substrate for initiation of the antiviral response and that their derivative antibodies were capable of neutralizing influenza and SARS-CoV-2.^68,69^ We found DENV-specific IgD+/IgM+ B cells persist, though they appear to diminish with multiple exposures. Further research should assess the role of these naïve-like B cells in the context of DENV infection.

Another key subset of MBCs with functional relevance are atypical MBCs, which have been previously implicated in deficient effector functions in the context of infection with HIV, hepatitis C virus, and *Plasmodium falciparum*.^29,71,72^ Interestingly, we detected consistently higher DENV-specific atypical MBCs after 2° compared to 1° dengue, suggesting a similar repeat-exposure-driven expansion of this cell subset in dengue. A prior study found high atypical MBCs in EC of DENV infection, and here we report that the DENV-specific subset of atypical MBCs is also high at EC.^37^ While we did not stimulate DENV-specific atypical MBCs to test their responsiveness, our hypothetical working model of DENV-specific B cell processes suggests that this subset may be responsive and give rise to subsequent DENV-specific activated and class-switched MBCs in 2° dengue. Accordingly, recent studies have also found that atypical MBCs can be responsive in the context of malaria.^25,73^ Understanding the impact of atypical MBCs on subsequent dengue immunity will be important.

Longitudinal evaluation of DENV-specific B cells from the same individuals over time provided two novel insights. First, the peak frequency of several DENV-specific B cell subsets occurred >3 months post-infection in 2° but not 1° DENV responses. Previously, Turner et al. demonstrated germinal center reactions up to ∼8 months post SARS-CoV-2 vaccination; thus, our finding of late peaks of DENV-specific MBCs is plausible.^74^ While germinal centers can give rise to incrementally more affinity-matured antibody specificities with time, there is heterogeneity across individuals in duration of germinal centers, similar to the heterogeneity in peak MBC response that we found.^75,76^ Secondly, we observed an unexpected uptick in frequencies of some DENV-specific B cell subsets between 12-18 months after 2° but not 1° dengue. While DENV-binding antibodies are thought to be boosted through seasonal exposure in endemic settings, it is unknown if DENV-specific MBCs can be similarly boosted.^77^ Previous studies indicate that DENV PB responses and seroconversion can occur in the absence of detectable viremia, suggesting that the virus-specific B cell compartment can be reshaped without disease.^78,79^ In sum, 2° dengue demonstrates distinct long-term MBC development processes from a 1° response, and this should be investigated further.

A key strength of our study is the detailed enumeration of multiple DENV-specific B cell subsets in the DENV-immune response in natural infections, including by isotype and MBC phenotype. To our knowledge, this is the first report classifying up to nine DENV-specific B cell subsets as well as two acute antibody-secreting cell responses from the same samples and individuals over time. Further, our longitudinal sampling approach from the same person enabled novel insights about distinct DENV-specific B cell dynamics in 1° versus 2° immunity. Finally, sampling up to 18 months post-infection provided an estimate of durability for DENV-specific B cell subsets. However, our study also has limitations, such as the small sample size of four participants in each 1° and 2° groups, which may result in an inability to detect some differences in DENV-specific subsets and low statistical power. Additionally, our convenience sampling design may limit generalizability. While our small sample size may have influenced the observed lack of significant differences in total frequency of DENV-specific B cells by infection history, this finding is corroborated by prior studies of total DENV-specific B cells.^33,37,80^ Finally, we evaluated symptomatic DENV1 and DENV3 serotype infections, and dynamics may differ for other DENV serotypes or milder inapparent infections.

Overall, our longitudinal enumeration of DENV-specific B cell compartments reveals that the 2° immunity consists not only of a quantitative boost, but a qualitative reprogramming of the memory pool. We found that DENV-specific B cell responses to 1° versus 2° dengue differ substantially in terms of subsets, despite no difference in total DENV-specific B cell frequency, suggesting an important role for phenotypically distinct B cell subsets in DENV immunity. We identified the presence and distinct kinetics of naïve-like and atypical DENV-specific B cells, as well as a potential role for DENV-specific IgM+ MBCs in initiating 2° immune responses. Intriguingly, we note that peak of DENV-specific MBCs occurs later than 3 months post-infection in 2° immunity, suggesting potential for long-term cellular processes. Harnessing these subset-specific and temporal differences will be critical for defining cellular correlates of protection and engineering next-generation vaccines capable of eliciting a superior memory pool.

## Materials and Methods

### Study Design

Longitudinal and Singlets pediatric PBMC samples were obtained from our ongoing Pediatric Dengue Hospital Study (PDHS) at the Hospital Infantil Manuel de Jesús Rivera (HIMJR), as previously described.^81^ Children aged 6 months to 14 years presenting with suspected DENV infection were enrolled. DENV infection was confirmed by RT-PCR in the acute phase and/or seroconversion by MAC-ELISA or inhibition ELISA (iELISA) and/or a ≥4-fold increase in the DENV iELISA titer between acute and early convalescence.^82–84^ For more detail, please see Supplementary Methods.

### Virus labelling with Alexa Fluor

Purified and/or concentrated DENV virions were conjugated to either AF488 or AF647 using an adaptation of a previously developed approach.^41,85^ Briefly, AlexaFluor succinimidyl ester (Invitrogen) was reconstituted in 0.2M sodium bicarbonate buffer (pH 8.3) and added to 200μL of purified DENV, or 500μL of concentrated DENV, at a final fluorophore concentration of 200uM or 100uM for AF488 and AF647, respectively. The mixture was incubated in the dark at room temperature for 1 hour with gentle inversion every 15 minutes. The labelling reaction was quenched with hour with gentle inversion every 15 minutes. Labeled DENV was purified by size exclusion chromatography on Sephadex G-25 columns (Cytiva), and fractions were collected, aliquoted, and stored at -80°C until further use. See Supplementary Methods for more details.

### Immunophenotyping of DENV-specific B cells

Cryopreserved PBMCs were rapidly thawed in a 37°C water bath, then resuspended in RPMI (Gibco) supplemented with 10% heat-inactivated fetal bovine serum (Corning) and 0.4nM Benzonase (Millipore). PBMCs were stained with Aqua Live/Dead viability dye (Thermo Fisher Scientific), CD3-V500 (clone UCHT1, Becton Dickenson [BD]), CD14-V500 (clone M5E2, BD), CD16-V500 (clone 3G8, BD), IgD-V450 (clone IA6-2, BD), IgM-BV605 (clone G20-127, BD), IgG-BV786 (clone G18-145, BD), CD19-APC-Cy7 (clone SJ25C1, BD), CD20-PerCP-Cy5.5 (clone 2H7, BD), CD21-PE-CF594 (clone B-ly4, BD), CD27-PE-Cy7 (clone O323, BD), and CD38-PE (clone HB7, BD) at 5μL per million cells as per the manufacturer’s specification, as well as optimized amounts of AF488-DENV1, AF488-DENV2, AF488-DENV3, AF647-DENV1, AF647-DENV2, and AF647-DENV3. Cells were stained and washed in 1% bovine serum albumin in PBS buffer supplemented with 5uM Chk2 kinase inhibitor II (Calbiochem) to prevent cell death. Stained cells were fixed with 4% paraformaldehyde in PBS for 15 minutes at room temperature, washed, and then resuspended in wash buffer for storage. Ultracomp eBeads (Thermo Fisher Scientific) were conjugated to each antibody used for cell staining (except for CD14-V500 and CD16-V500), as well as CD32-AF647 (clone FUN-2, Biolegend) and CD95-AF488 (clone DX2, Biolegend) according to manufacturer’s instructions for use as single-color compensation controls prior to every data collection. Cells were analyzed with a BD LSRFortessa (BD), and data were analyzed in FlowJo. The boundaries for the CD21- and CD27-positive cell populations were determined through fluorescence minus one control (FMO) and acquired for each experiment. To control for autofluorescence, we used an unstained PBMC sample in each experiment. The negative control for DENV-specific B cells was a PBMC sample stained in the same manner, but without any AF-labelled DENV antigens. No-antigen control was acquired in each experiment and used to define the negative population in establishing the threshold for DENV-positivity. For the longitudinal sample set, each individual’s set of six longitudinal samples from acute to 18M was stained on the same day and acquired in one experiment to minimize batch effect for temporal analyses.

## Acknowledgements

We are deeply grateful to the participants of the Pediatric Dengue Hospital Study (PDHS) and their families for their continued commitment. We greatly appreciate the dedication and invaluable contributions of the past and present study team members at the Hospital Infantil Manuel de Jesús Rivera (HIMJR), the Sustainable Sciences Institute in Nicaragua, and the Laboratorio Nacional de Virología at the Centro Nacional de Diagnóstico y Referencia in Managua. In particular, we are grateful to Raul Zapata and others for excellent PBMC sample processing and cryopreservation, and to the Nicaraguan clinical and field teams who followed up with participants for longitudinal sampling. We also appreciate the work of Carlos Montenegro and Jorge Ruiz for data management and collating participant datasets. Importantly, we thank Dr. Claudia Sanchez San Martin, Erika Vanessa Posada, Marco Chapa, and Agamjot Bal at the University of California, Berkeley, for inventory of liquid nitrogen tanks and supporting PBMC sample selection. Finally, we are grateful to Dr. Mark Slifka at Oregon Health & Science University for reviewing the manuscript and providing helpful comments.

## Funding Sources

This work was supported by grant P01 AI106695 (EH) from the National Institute for Allergy and Infectious Diseases of the National Institutes of Health (NIAID/NIH). The PHDS was supported by NIAID/NIH grants R42 AI062100 (EH), R01 AI099631 (AB) and U54 AI065359 (AB; Program Director Alan Barbour). During this project, T.S. was supported by the HHMI Hanna H. Gray Fellowship (Grant ID: GT16780) and a small grant from the UC Berkeley Center for Emerging and Neglected Diseases, and A.B. was supported by the UC Berkeley Summer Undergraduate Research Fellowship.

## Data Sharing Plan

The anonymized data for this study are included as a supplementary appendix upon publication. The custom R code will be available on GitHub upon publication. Please address any other correspondence and/or data requests to Eva Harris.

## Notes

### Competing Interest Statement

The authors have declared no competing interest.

