## Supplemental for "Dengue virus-specific memory B cell subsets differ as a function of infection history"

This PDF includes

- Supplementary Methods (pg. 2 – 4)
- Supplementary Tables (pg. 5 – 8)
- Supplementary Figures (pg. 9 – 25)

### Supplementary Methods

**Details on study design.** Blood samples were collected upon presentation to the hospital (acute phase) and at follow-up visits during early convalescence (~14–28 days post-symptom onset), 3 months, 6 months, 12 months, and 18 months post-symptom onset. Samples for this study were selected from subjects with symptomatic dengue fever, confirmed DENV infection, and evidence of a productive immune response, defined by a ≥4-fold increase in DENV iELISA titers between enrollment and early convalescence. DENV infection history was classified based on total anti-DENV antibody titers measured by inhibition ELISA (iELISA) in paired acute- and convalescent-phase samples.[1] Specifically, primary DENV infection was defined by an iELISA titer <2560 in the early convalescent phase, while a secondary infection was indicated by an iELISA titer of ≥2560 in the convalescent phase. The naïve pediatric PBMC sample was obtained from the companion Pediatric Dengue Cohort Study (PDCS), an ongoing prospective open cohort study of ~4,000 children (ages 2–17) in Managua, Nicaragua.[2] Naïve status was defined by a negative result by DENV iELISA at enrollment and negative during follow-up.[1] PBMC samples that were used as the no-antigen negative control were obtained from anonymous adult blood donations to the Centro Nacional de Sangre in Managua, Nicaragua.

**Sample collection and PBMC preparation.** Whole blood was collected in EDTA Vacutainer tubes (Becton Dickinson). Upon receipt at the Nicaraguan National Virology Laboratory—within 2 hours of collection to ensure sample integrity—4–5 mL of the EDTA-treated blood was layered onto Leucosep tubes (Greiner Bio-One) preloaded with 3 mL of Ficoll Histopaque (Sigma) and centrifuged at 500×g for 20 minutes at room temperature. The PBMC layer was transferred to a 15-mL conical tube containing 9 mL of PBS supplemented with 2% FBS (Denville Scientific) and 1% penicillin-streptomycin (Sigma), washed three times (500×g for 10 minutes each). Before the final wash, an aliquot was removed for cell counting using a Sysmex XS-1000i hematology analyzer. After the last wash, cells were adjusted to $1\times10^7$ cells/mL in freezing medium (90% FBS, 10% DMSO), aliquoted into cryovials, cooled in isopropanol-based freezing containers (Mr. Frosty; Nalgene) at −80°C overnight, and then transferred to liquid nitrogen for long-term storage. All steps—from blood draw through cryopreservation—were time-stamped to maintain cold-chain and quality control. Finally, PBMC samples were shipped to the UC Berkeley Harris Laboratory in liquid-nitrogen dry shippers for downstream analyses.

**Ethics statement.** The human subjects protocols for the PDHS and the PDCS were reviewed and approved by the Institutional Review Boards of the University of California, Berkeley (2010-06-1649 and 2010-09-2245, respectively), and the Nicaraguan Ministry of Health (NIC-MINSA/CNDR CIRE-01/10/06-13.Ver7 and NIC-MINSA/CNDR CIRE-09/03/07-008.ver10, respectively). Parents or legal guardians of all participants provided written informed consent, and participants 6 years of age and older provided assent.

**Virus production.** Vero-furin cells (kindly provided by Dr. Ralph Baric at the University of North Carolina, Chapel Hill) were cultured in growth medium consisting of DMEM (Gibco), 1x non-essential amino acids (Gibco), 1x sodium pyruvate (Gibco), 1x penicillin-streptomycin (Gibco), and 5% heat-inactivated fetal bovine serum (Corning) at 37°C and 5% $CO_2$. Nicaraguan DENV strains were isolated from clinical samples and subsequently propagated in Vero-furin cells using an MOI of 0.025 to increase the level of maturation of the virion in order to more closely resemble those of the virion *in vivo* during human infection.[3,4] Viruses were produced in multiple flasks to generate large batches of the same stock. Cell supernatants containing viruses (~250mL) were collected 5-6 days post-inoculation, centrifuged to pellet cell debris, filtered, and then concentrated ≥10-fold in 70-mL centrifugal spin filters (Millipore Sigma) prior to storage at -80°C. Concentrated DENV was purified by ultracentrifugation using an 30% Optiprep sucrose gradient at 30,000xg for 3.33 hours in a vacuum at 4°C, without brakes. Purified DENV was collected from the interface of the 30% and 55% Optiprep gradient with a syringe and then stored at -80°C until labeling.[5]

**Bead-based confirmation of virus labelling.** To determine the optimal fraction of labelled DENV and confirm labelling of DENV, binding to a mAb-conjugated bead was assessed via flow cytometry. One drop of UltraComp eBeads (Invitrogen) was dispensed in flow tubes, then conjugated according to the manufacturer's instructions to DENV-binding serotype-specific monoclonal antibodies (mAbs) that bind a quaternary epitope on the virion surface: 14C10, 2D22, and 5J7, to target DENV1, DENV2, and DENV3, respectively (kind gift from Ralph Baric and Aravinda de Silva at the University of North Carolina, Chapel Hill). Beads were subsequently washed with a PBS/BSA buffer (1x PBS buffer and 1% bovine serum albumin), incubated with optimized dilutions of the corresponding serotype of labelled virus antigen for 30 minutes at 4°C, fixed with 4% paraformaldehyde for 15 minutes at RT, washed, and then resuspended in PBS/BSA. Beads were analyzed on the LSRFortessa (Becton Dickenson), with a minimum of 10,000 events acquired per sample and then analyzed with FlowJo. CD95-AF488 (clone DX2, Biolegend) and CD32-AF647 (clone FUN-2, Biolegend) antibodies directly conjugated to the beads were used as positive controls for AF488 and AF647 signal, respectively, and unstained beads (i.e., no antibody) were used as the negative control.

**Quantification of DENV particles in labelled antigens.** Each labelled antigen (100µL) was added to 40µL of molecular biology grade water (Corning), and viral RNA was extracted as per the manufacturer's instructions using the QIAmp Viral RNA Kit (Qiagen). The final RNA elution was performed using two elutions of 30µL, each using molecular biology grade water with a spin of 6000xg for 1 minute. RNA was stored in 5-µL aliquots at -20°C. Subsequently, viral particles were quantified as described previously.[3] Briefly, extracted RNA was reverse-transcribed to cDNA as per the manufacturer's instructions using the GoTaq 2-step RT-qPCR System (Promega). Quantification was performed against a standard curve of an amplicon containing cDNA encompassing nucleotides 10361-10660 corresponding to the 3' untranslated region of DENV2 clinical isolate NI15. Pan-DENV primers (forward: TTGAGTAAACYRTGCTGCCTGTAGCTC, reverse: GAGACAGCAGGATCTCTGGTCTYTC) were utilized. The quantity of viral particles was calculated in units of cDNA equivalents per mL.

**Data analysis.** Cell gating was performed manually in FlowJo (version 10.10.0), and cell frequencies were exported and organized using Microsoft Excel. Data was visualized and analyzed using FlowJo, GraphPad Prism (version 10), and RStudio (version 4.4.2; Packages: tidyverse, readxl, rstatix, ggplot2, ggtext, ggprism, gtools, Hmisc, corrplot, pheatmap, igraph, ggraph, scales, grid, graphlayouts, ggpubr, dplyr, and showtext). This approach minimizes total within-cluster variance. *Cross-sectional differences* in the 1° versus 2° DENV-specific B cell frequencies (at single timepoints) were tested via unpaired two-sided or one-sided Wilcoxon Rank Sum test. The one-sided test was specifically performed to test the hypothesis that values in the 1° infection group were less than those in the 2° infection group at each specific timepoint. The alpha threshold for significance was set to 0.05 and not adjusted for multiple comparisons. *Differences in the longitudinal dynamics* of DENV-specific B cells and overall B cell populations were tested using Two-Way Repeated Measures Analysis of Variance (ANOVA). Specifically, the overall effect of infection history (between-subjects factor) and time (within-subject factor) was analyzed, grouping by participant, with an alpha threshold for significance set to 0.05. The *Directed Network Graph* enabled visualization of correlations in DENV-specific B cell or overall B cell frequency over sequential time points (i.e., Acute → EC, EC → 3M, 3M → 6M, 6M → 12M, and 12M → 18M). Pairwise correlations of (1) DENV-specific cells per 10,000 parental B cells and (2) percent B cell subset out of total B cells were conducted using the Spearman Rank test. The alpha level to define significance in these correlations was set to ≤0.1, and only significant correlations were plotted. Also, positive correlations (*rho*>0) were selected. For some plots, known biologically implausible cellular transitions (i.e., IgD-/IgG+ → IgD+) were filtered out as per the criteria in Table S4 to visualize correlations that may be biologically plausible. Distinct B cell subsets were plotted as colored nodes, and all significant correlations were plotted as an arrow from the earlier timepoint directed to the later timepoint. The thickness of each arrow corresponded to the absolute magnitude of *rho*. This resulting network was visualized using a Sugiyama layout for directed acyclic graphs which enabled creation of distinct layers in the plot based on timepoint to depict the longitudinal structure of the underlying data.

### Supplementary Tables

**Table S1. Participant characteristics of longitudinal sample set up to 18 months post-infection[a].**

| Infection History | Study ID | Age | Sex | iELISA acute | iELISA EC | DENV serotype | Severity[b] | Year of Infection |
|---|---|---|---|---|---|---|---|---|
| Primary | P1 | 9 | M | <10 | 588 | 1 | Severe | 2012 |
| Primary | P2 | 11 | M | 12 | 460 | 3 | DwoWS | 2011 |
| Primary | P3 | 11 | F | <10 | 136 | 3 | DwWS | 2011 |
| Primary | P4 | 6 | M | <10 | 374 | 1 | DwWS | 2012 |
| Secondary | S1 | 12 | F | 2819 | 25968 | 1 | DwoWS | 2011 |
| Secondary | S2 | 9 | M | 1274 | >100000 | 1 | Severe | 2011 |
| Secondary | S3 | 11 | M | 3384 | 49292 | 3 | Severe | 2011 |
| Secondary | S4 | 14 | M | 6498 | >100000 | 3 | DwWS | 2011 |

[a]All pediatric cases are from the PDHS with RT-PCR-confirmed DENV infection.
[b]Disease severity as per WHO 2009 Guidelines: Dengue without Warning Signs (DwoWS), Dengue with Warning Signs (DwWS), and Severe Dengue (Severe).

**Table S2. Participant characteristics of early single timepoint sample set[a].**

| Infection History | Study ID | Age | Sex | iELISA Acute | iELISA EC | DENV serotype | Severity[b] | Year of Infection | Timepoint |
|---|---|---|---|---|---|---|---|---|---|
| Primary | P5 | 11 | M | 124 | 163 | 2 | DwWS | 2005 | EC |
| Primary | P6 | 12 | F | 20 | 320 | 2 | DwWS | 2005 | EC |
| Primary | P7 | 7 | M | <10 | 125 | 3 | DwWS | 2007 | 3M |
| Primary | P8 | 5 | F | <10 | 23 | NA | Severe | 2010 | EC |
| Primary | P9 | 9 | M | <10 | 12 | 3 | DwWS | 2010 | EC |
| Secondary | S5 | 12 | F | 128 | 30994 | 2 | DwWS | 2015 | EC |
| Secondary | S6 | 7 | M | <10 | 24115 | 2 | DwWS | 2007 | 3M |
| Secondary | S7 | 14 | F | 24997 | 81063 | 3 | DwWS | 2007 | 3M |
| Secondary | S8 | 12 | M | 137 | 27856 | 3 | DwWS | 2010 | EC |
| Secondary | S9 | 12 | M | 196 | >100000 | 2 | DwWS | 2008 | 3M |

[a]All pediatric dengue cases are from the PDHS with RT-PCR-confirmed DENV infection.
[b]Disease severity as per WHO 2009 Guidelines: Dengue without Warning Signs (DwoWS), Dengue with Warning Signs (DwWS), and Severe Dengue (Severe).

**Table S3. Definitions of B cell subsets from parental gate (CD3-/CD14-/CD16-/ LIVE+).**

| Cell type | Definition |
|---|---|
| **Naïve and IgD+ subsets** | |
| IgD+ naïve | CD19+/ CD20+/ IgD+ |
| IgD+/IgM+ cells | CD19+/ CD20+/ IgD+/ IgM+ |
| **Antibody Secreting Cells** | |
| PB/PC<br>*(Plasmablasts and plasma cells)* | CD19+/ CD20$^{-/low}$/ IgD-/ CD27+/ CD38+ |
| IgM- PB<br>*(estimates largely IgG PBs and some IgA PBs. Note that IgG PBs express variable B cell surface receptor)* | CD19+/ CD20$^{-/low}$/ IgD-/ CD27+/ CD38+/ IgM- |
| IgM+ PB | CD19+/ CD20$^{-/low}$/ IgD-/ CD27+/ CD38+/ IgM+ |
| **Memory B cells (MBCs) and IgD- subsets** | |
| Class-switched IgD- MBC | CD19+/ CD20+/ IgD- |
| IgG+ MBC | CD19+/ CD20+/ IgD-/ IgM-/ IgG+ |
| IgM+ MBC | CD19+/ CD20+/ IgD-/ IgM+ |
| IgD-/IgM-/IgG- MBC<br>*(estimates largely IgA MBCs in periphery)* | CD19+/ CD20+/ IgD-/ IgM-/ IgG- |
| Resting MBC | CD19+/ CD20+/ IgD-/ CD27+/ CD21+ |
| Activated MBC | CD19+/ CD20+/ IgD-/ CD27+/ CD21- |
| Atypical MBC<br>*(This subset is inclusive of atypical MBCs)* | CD19+/ CD20+/ IgD-/ CD27-/ CD21- |

**Table S4. Exclusion criteria for biologically implausible cellular transitions among direct correlations of DENV-specific B cell frequency over consecutive timepoints.**

| Exclusion criteria | Reasoning |
|---|---|
| Exclude if any cell type from the first timepoint is correlated with IgD+ naïve or IgD+/IgM+ B cells in the second timepoint.<br><br>*Examples of excluded correlations:*<br>Resting memory B cell → IgD+ naïve<br>Resting memory B cell → IgD+/IgM+<br>IgD-/IgM-/IgG- → IgD+ naïve<br>IgD-/IgM-/IgG- → IgD+/IgM+<br>IgD-/IgM+ → IgD+ naïve | Due to order of isotype switching based on irreversible splicing events, IgD- B cell subsets cannot revert to expressing IgD. |
| Exclude if IgG+ or IgD-/IgM-/IgG- B cells are correlated from one timepoint are correlated with IgM+ B cells in the next timepoint<br><br>*Examples of excluded correlations:*<br>IgG+ → IgM+<br>IgD-/IgM-/IgG- → IgM+ | Due to order of isotype switching based on irreversible splicing events, IgG+ and IgD-/IgM-/IgG- B cells cannot revert to expressing IgM. |

### Supplementary Figures

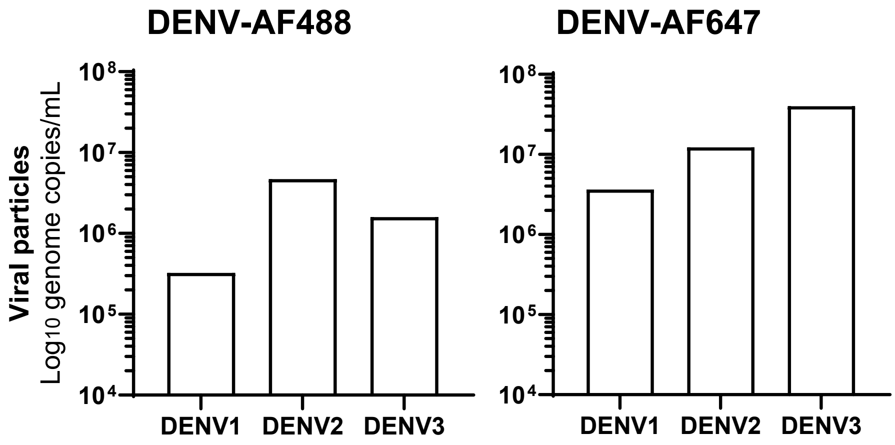


**Fig S1. Presence of viral particles in fluorescently labelled DENV antigen preparations.** DENV1, DENV2, and DENV3 viruses were labelled with either AF488 or AF647 and purified via size exclusion chromatography. Viral particles per mL were quantified via RT-qPCR in the optimal fraction used to stain B cells. Viral titer was interpolated from a known cDNA standard curve.

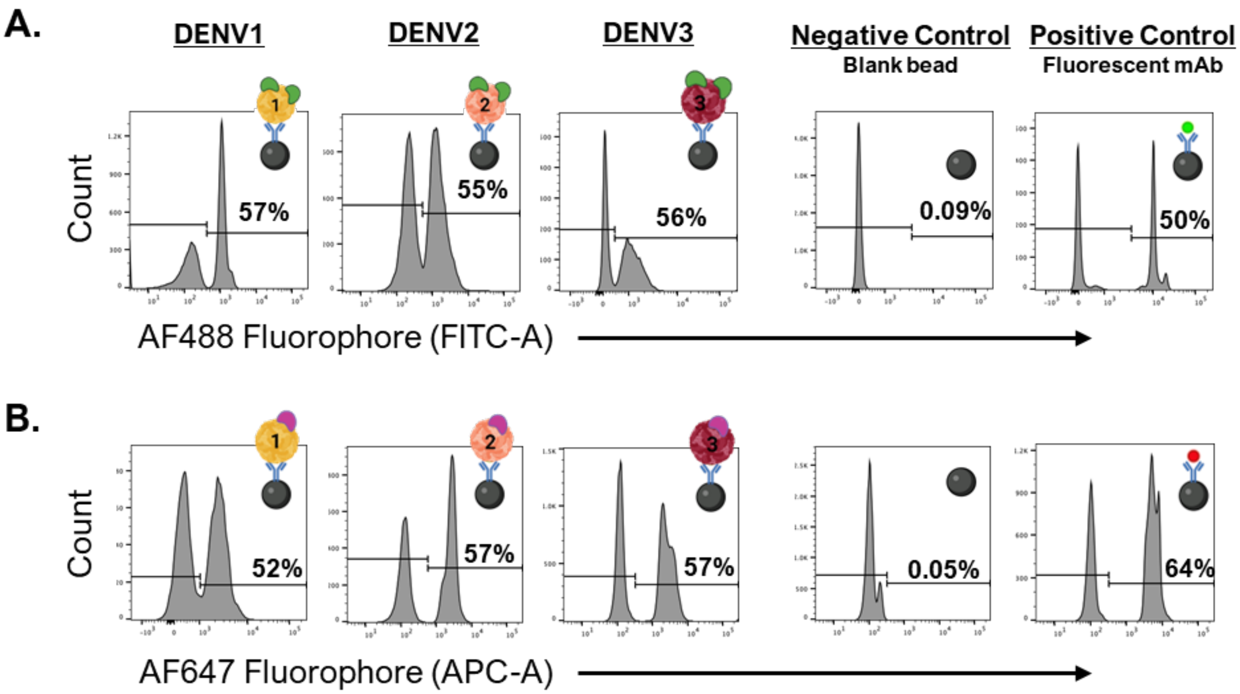


**Fig S2. Representative quality control to confirm successful labelling of virus particles.** DENV1, DENV2, and DENV3 viruses were labelled with either AF488 (**A**) or AF647 (**B**). Each labelled preparation was co-incubated with Ultracomp beads bound to DENV-specific monoclonal antibodies (mAbs). DENV-specific mAbs used in this approach each target a serotype-specific quaternary epitope: 14C10 mAb against DENV1; 2D22 mAb against DENV2; 5J7 mAb against DENV3. If the mAb-bound Ultracomp bead captured fluorescently tagged antigen, a positive fluorescence peak was detectable in either the FITC channel for AF488 fluor or the APC channel for AF647 fluor. Thus, a positive peak confirmed the presence of labelled DENV and the availability of a type-specific quaternary epitope on the fluorescently tagged DENV antigens. Ultracomp bead counts are shown by Mean Fluorescence Intensity. The percent of beads with a positive threshold of fluorescence is indicated. The negative control is the blank bead, and the positive control is the bead conjugated with an AF488 or AF647 fluorescently labelled mAb.

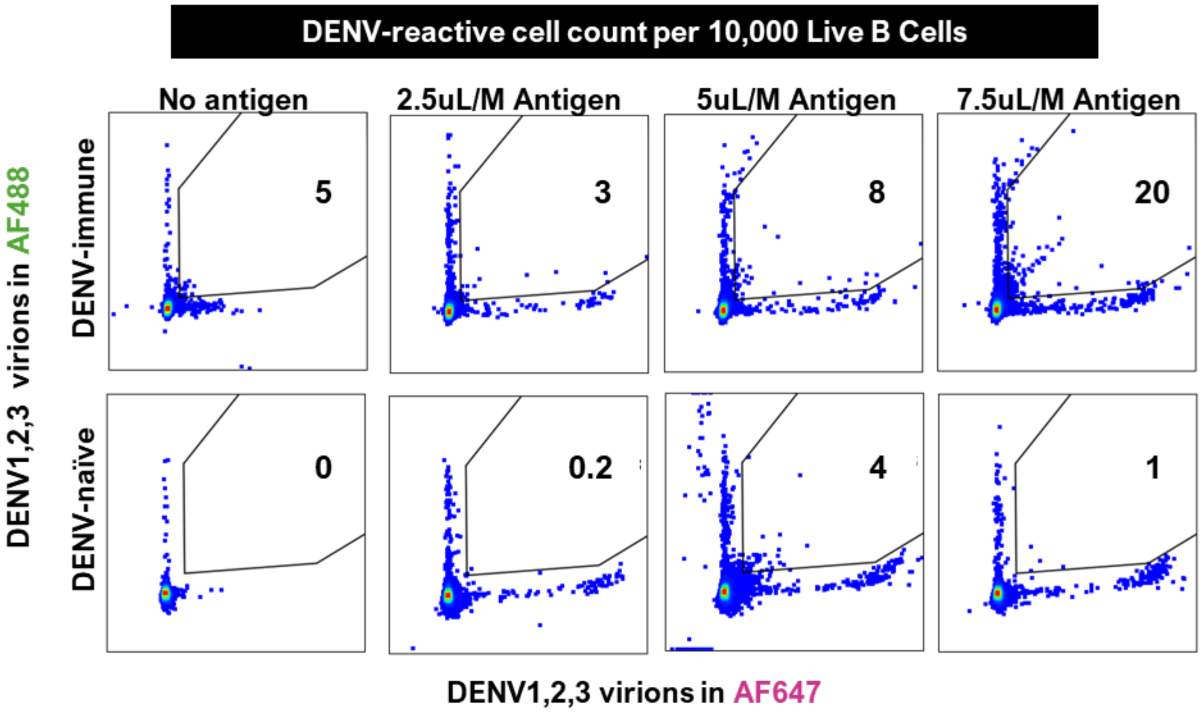


**Fig S3. Titration of antigen cocktail on DENV-immune versus DENV-naïve case.** DENV1, DENV2, and DENV3 viruses were each labelled with AF488 or AF647, generating six single-label viruses that were combined in equal volumes to develop an antigen cocktail. This cocktail was well-mixed and titrated by microliter volume per million PBMCs (uL/M) on a DENV-immune sample from the early convalescence of a secondary DENV2 infection, as well as a DENV-naïve sample. The DENV-naïve sample was derived from the Nicaraguan Pediatric Dengue Cohort Study, and DENV-naïve serostatus was defined based on no detectable DENV-binding antibodies in the DENV iELISA. From total live B cells (Singlets/Live/CD3-/CD14-/CD16-/CD19+), the AF488 and AF647 double-positive gate is shown, with the number of DENV-specific B cells counted per 10,000 parental B cells. Antigen dose-dependent increase is observed in the DENV-immune sample, and the artefactual double-positive cells are substantially lower in the DENV-naïve sample, indicating that DENV-reactive B cells can be specifically detected in this system.

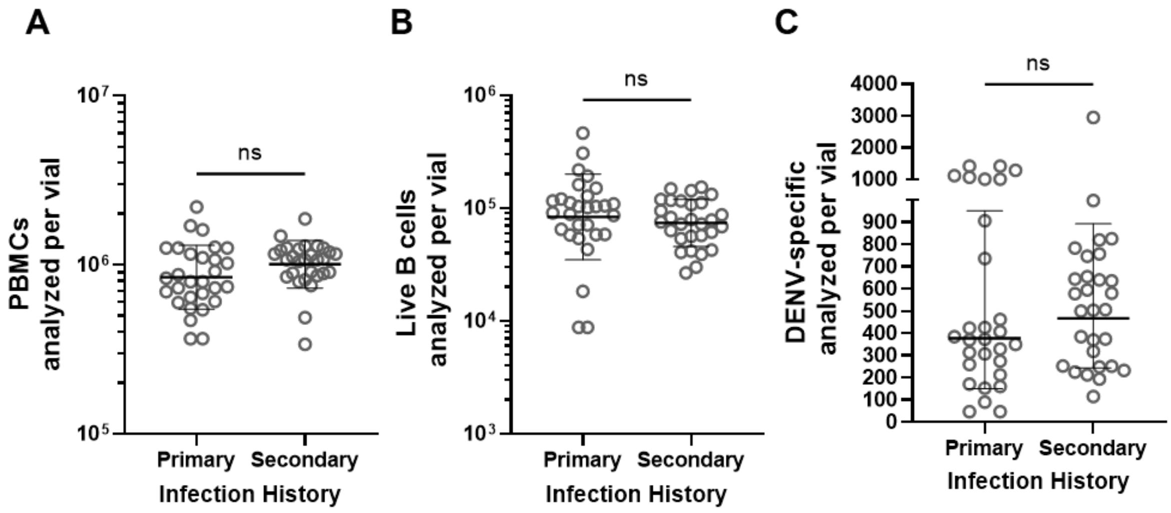


**Fig S4. Summary of raw cell counts analyzed in this study.** A total of 58 PBMC samples (i.e., cryovials) collected from pediatric dengue cases were analyzed in this study as part of the Longitudinal and Singlets sample sets. Per-sample counts of PBMCs (**A**), live B cells (**B**), and DENV-reactive B cells (**C**) are indicated. Differences in raw counts by primary versus secondary DENV infection history are shown, assessed using t-test with line at the mean with standard deviation, with non-significance indicated (ns).

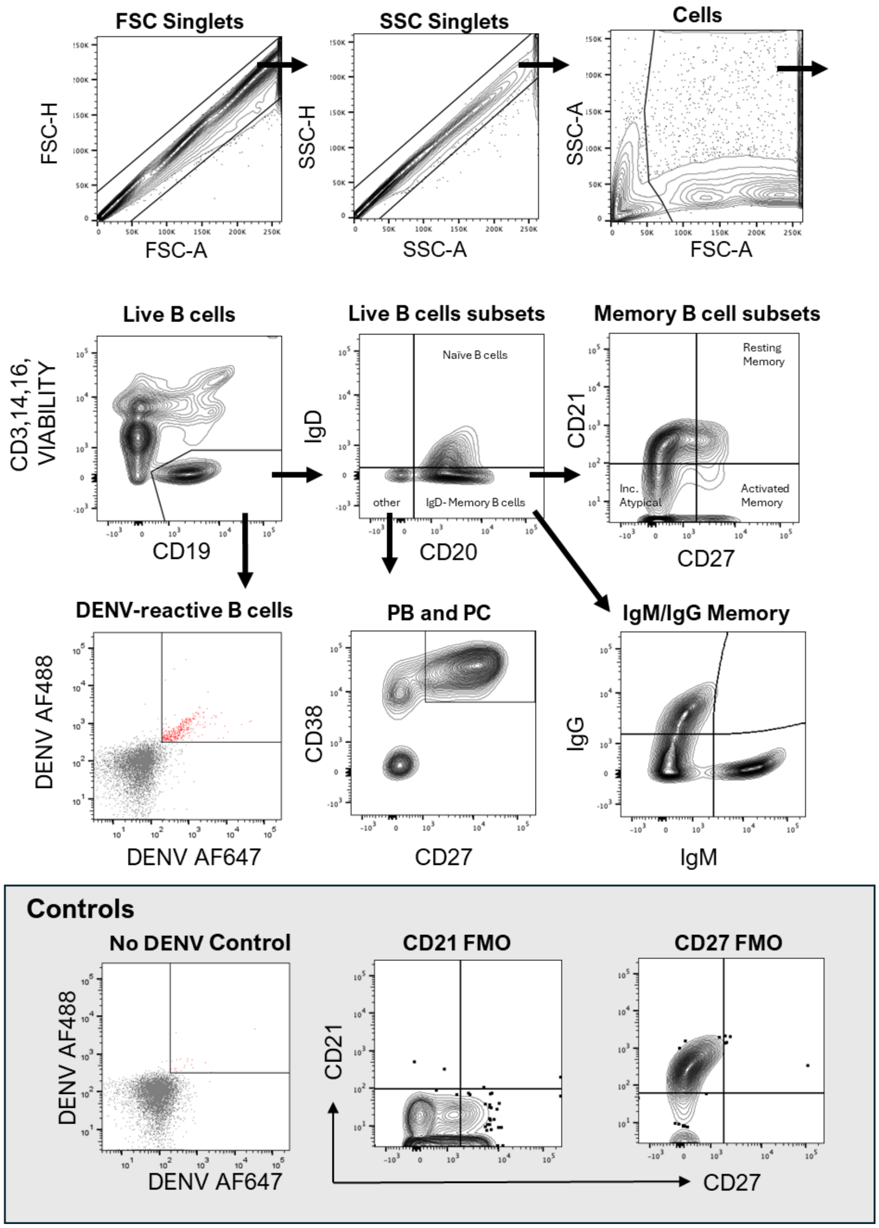


**Fig S5. Representative flow cytometry gating strategy of B cell subsets and controls.** Singlets were filtered based on forward scatter (FSC) and side scatter (SSC), and then debris was eliminated. From this, live B cells were gated as CD3-/CD16-/CD14-/AquaVitalDye- (all in same channel for dump), and CD19+. From total live B cells, naïve B cells and class-switched IgD- MBCs were defined. Class-switched IgD- MBCs were gated by CD21 and CD27, which enabled definition of resting MBCs, activated MBCs, and a double-negative population that includes atypical MBCs. Class-switched IgD- MBCs were also gated by isotype to IgG+, IgM+, and IgG-/IgM- (inferred IgA) MBCs. From each of these B cell subsets, a DENV-reactive gate was defined as AF488+/AF647+. Plasmablasts and plasma cells (PB and PC) were defined from the IgD-/CD20- subset as CD38+/CD27+. Fluorescence minus one (FMO) controls were used to define the threshold for positivity for CD27 and CD21. A PBMC sample without any DENV antigen, but all the cell surface marker fluorescent antibodies, was used to develop the gate for DENV-specific B cells and to evaluate the limit of detection.

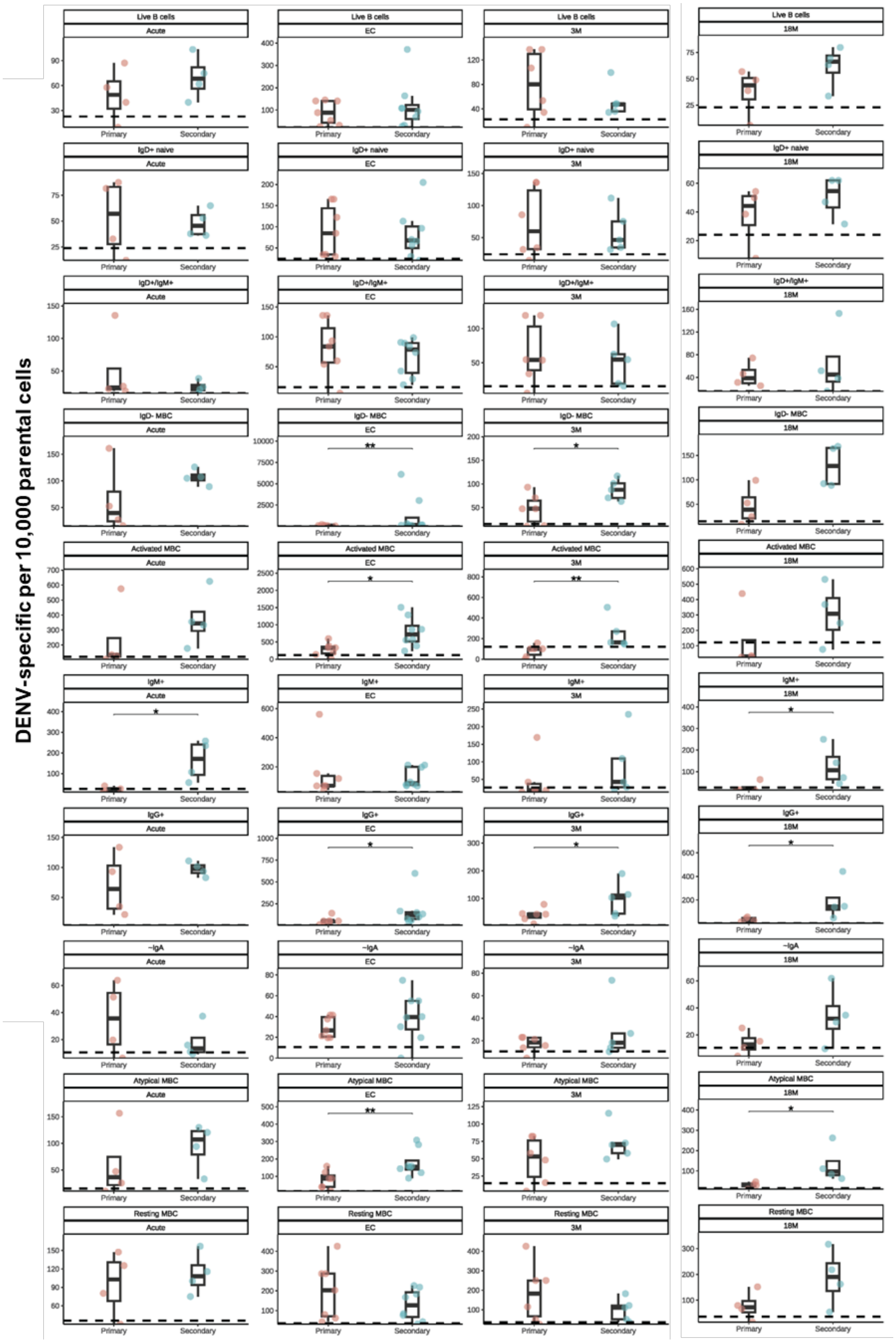
DENV-specific per 10,000 parental cells
Live B cells
IgD+ naive
IgD+/IgM+
IgD- MBC
Activated MBC
IgM+
IgG+
-IgA
Atypical MBC
Resting MBC
Acute
EC
3M
18M
Primary
Secondary

**Fig S6. Differences in DENV-specific B cell frequencies at each timepoint.** Frequency of DENV-specific B cells per 10,000 parental cells for each subset (rows) by 1° (pink) and 2° (blue) dengue groups over time (columns). Box plot with median and range is overlaid. Significant differences were assessed by one-sided Mann Whitney tests, where (*) indicates a $p<0.05$ and (**) indicates a $p<0.01$, with the hypothesis that the frequency of DENV-specific B cells will be higher in 2° than 1° dengue cases. Non-significant differences are indicated with blank. Dotted line indicates limit of detection based on mean of PBMCs without antigen for each subset (n=7).

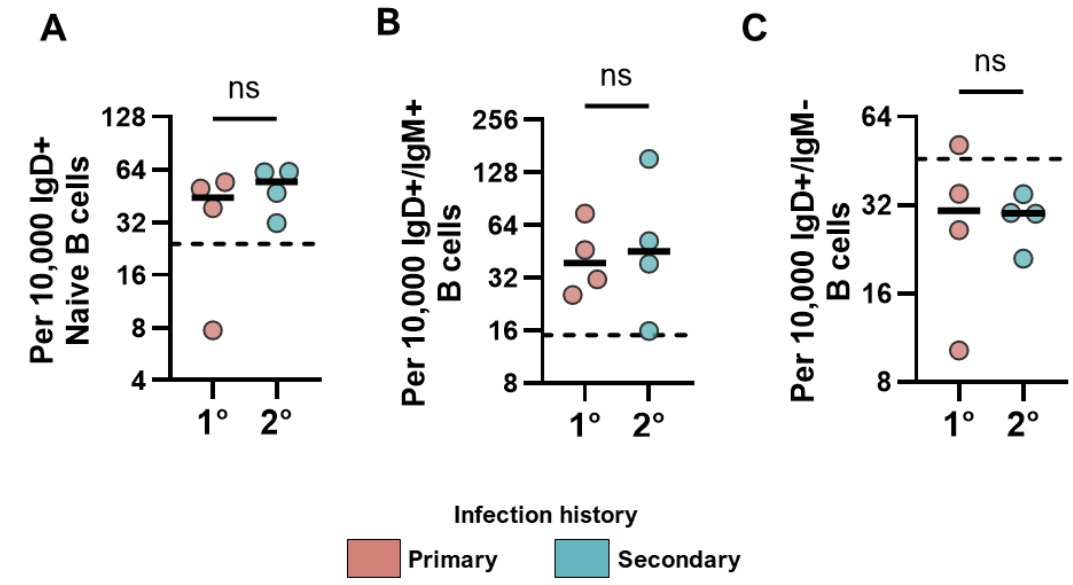


**Fig S7. Naive-like DENV-reactive cells at 18 months post-infection.** From the 18M samples in the Longitudinal Set, (**A**) naïve B cells were gated as (CD3-/CD16-/CD14-/CD19+/CD20+/IgD+), and within this, (**B**) IgD+/IgM+ double-positive cells and (**C**) IgD+/IgM- single-positive cells were gated. The dotted line indicates the limit of detection based on mean of PBMCs without antigen for each subset (n=7), showing that the IgD+/IgM- single-positive cells are largely artefactual. Primary dengue cases (1° n=4 participants) are shown in pink and secondary dengue cases (2° n=4 participants) are shown in blue. One-tailed Mann-Whitney test was used to test for significant differences, and the resulting non-significance (ns) is indicated on the plot.

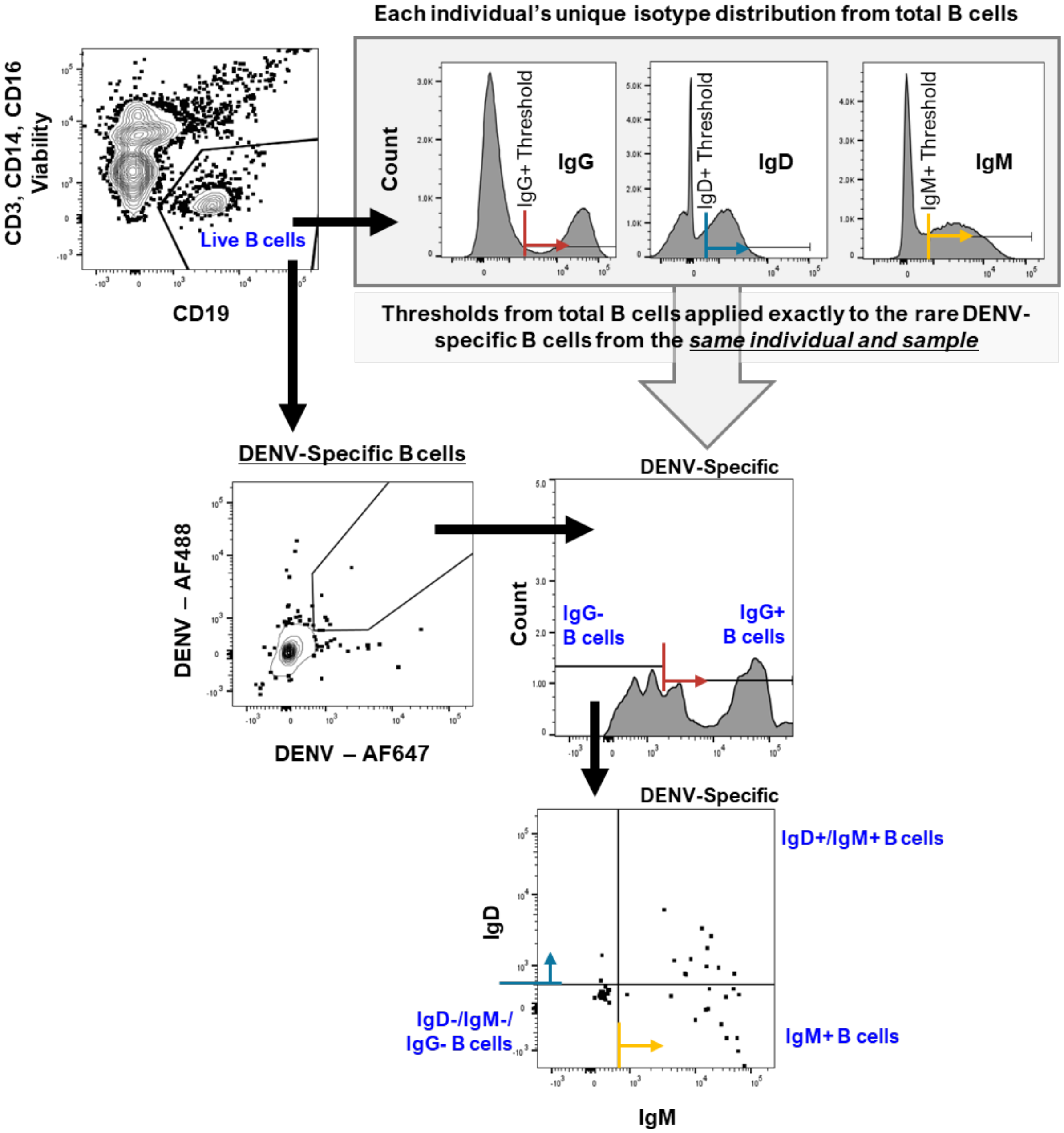


**Fig S8. Representative second gating strategy to assess isotype distribution among DENV-specific B cells.** Longitudinal Set PBMC samples from the 18M timepoint of the 1° (n=4) versus 2° (n=4) dengue groups were further evaluated to tailor isotype cut-off thresholds to each individual's phenotype and re-assess the extent of naïve-like DENV-specific cells (IgD+/IgM+). From well-sampled live B cells (CD3-/CD16-/CD14-/CD19+), the distribution of mean fluorescence intensity for each BCR isotype was derived in order to generate individual-specific limits of isotype-positivity thresholds (grey inset box). Colored lines and arrows indicate unique positivity thresholds for IgG+ (red), IgD+ (blue) and IgM+ (yellow) B cells. Unique positivity thresholds were determined per individual and sample, and the same positivity threshold was then applied to the sparse DENV-specific B cell population. Among a selection of bright subset of DENV-reactive B cells (AF488+/AF647+), the IgG+ and IgG- B cells were defined. From this IgG- B cell population, IgD-/IgM+, IgD+/IgM+, and IgD-/IgM- (inferred IgA) B cells were defined. This approach yielded the percentage of DENV-specific B cells occupied by mutually exclusive BCR isotypes.

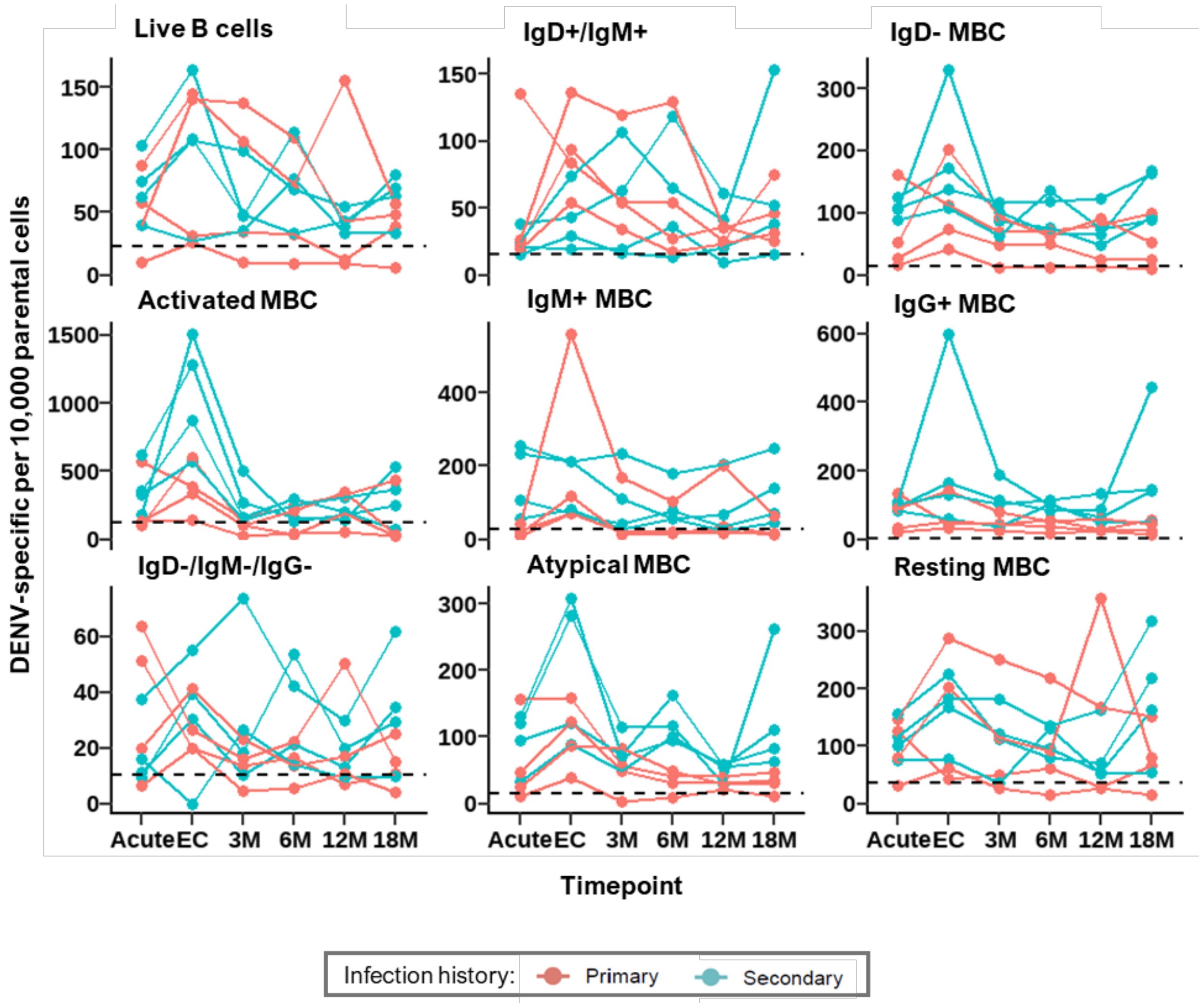


**Fig S9. Longitudinal trajectory of DENV-specific B cell frequency <u>per 10,000</u> parental cells over time.** Analytic dataset of longitudinal DENV-specific B cell frequencies per 10,000 parental cells for each of the nine key B cell subsets evaluated in this study. A frequency per 10,000 of parental cells is a <u>relative</u> metric that allows for comparisons within a subset over time. Each solid line represents data from a single participant from either the 1° (n=4 participants; pink) or 2° (n=4 participants; blue) dengue group. Participants were sampled over time at acute, EC, 3M, 6M, 12M and 18M. DENV-specific B cell subsets abbreviated as follows: IgD-/IgG-/IgM- (inferred IgA), IgD- MBC (class-switched MBCs), IgG+ MBC (IgD-/IgG+), IgM+ MBC (IgD-/IgM+) and IgD+/IgM+ (naïve-like B cells). Dotted line indicates limit of detection based on mean PBMC samples without antigen (n=7).

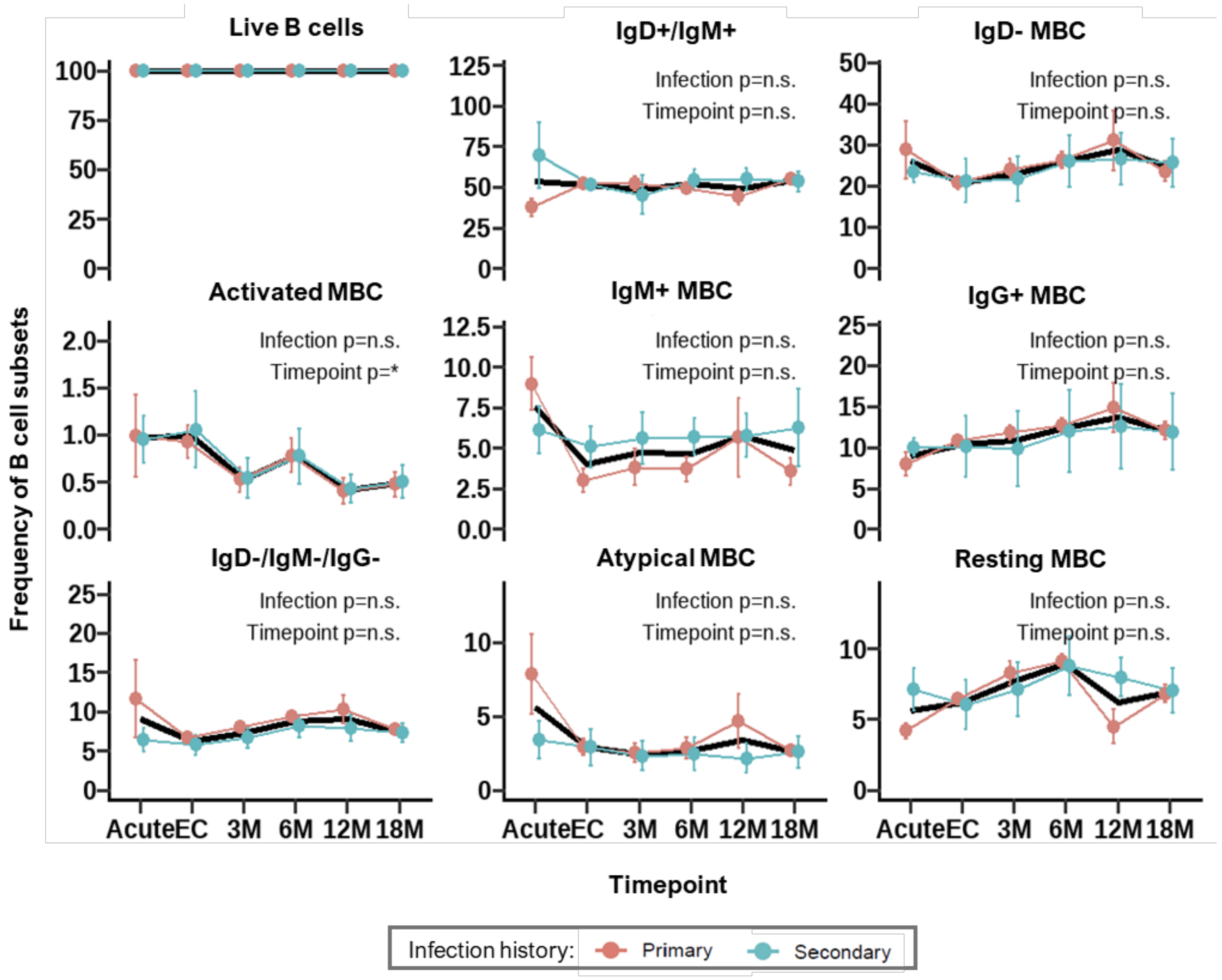


**Fig S10. No significant differences observed in <u>total</u> B cell subset frequency by DENV infection history.** To assess whether the distinct dynamics observed in the DENV-specific B cell compartment were merely an attribute of temporal changes in frequency of total B cells (not just the DENV-specific B cells), the differences in B cell subset frequency by infection and timepoint were tested. Two-way Repeated Measures ANOVA was conducted to evaluate the main effect of timepoint and infection history on B cell frequency in the 1° (n=4 individuals and n=6 timepoints; pink) versus 2° (n=4 individuals and n=6 timepoints; blue) infection history groups, accounting for individual variability. Timepoint was treated as a within-subject repeated measure, and infection history group (i.e., infection) was treated as a between-subjects measure. Mean frequency of B cell subsets as a percentage of live B cells was evaluated at each timepoint (Acute, EC, 3M, 6M, 12M, and 18M) and shown with standard error of the mean for each subset. The results of the test are annotated on each panel, showing the main effect significance for Infection and Timepoint, with non-significant differences indicated with "n.s." and a significant difference of $p<0.05$ indicated with (*). The solid black line represents the main effect of timepoint, which combines 1° and 2° infection groups.

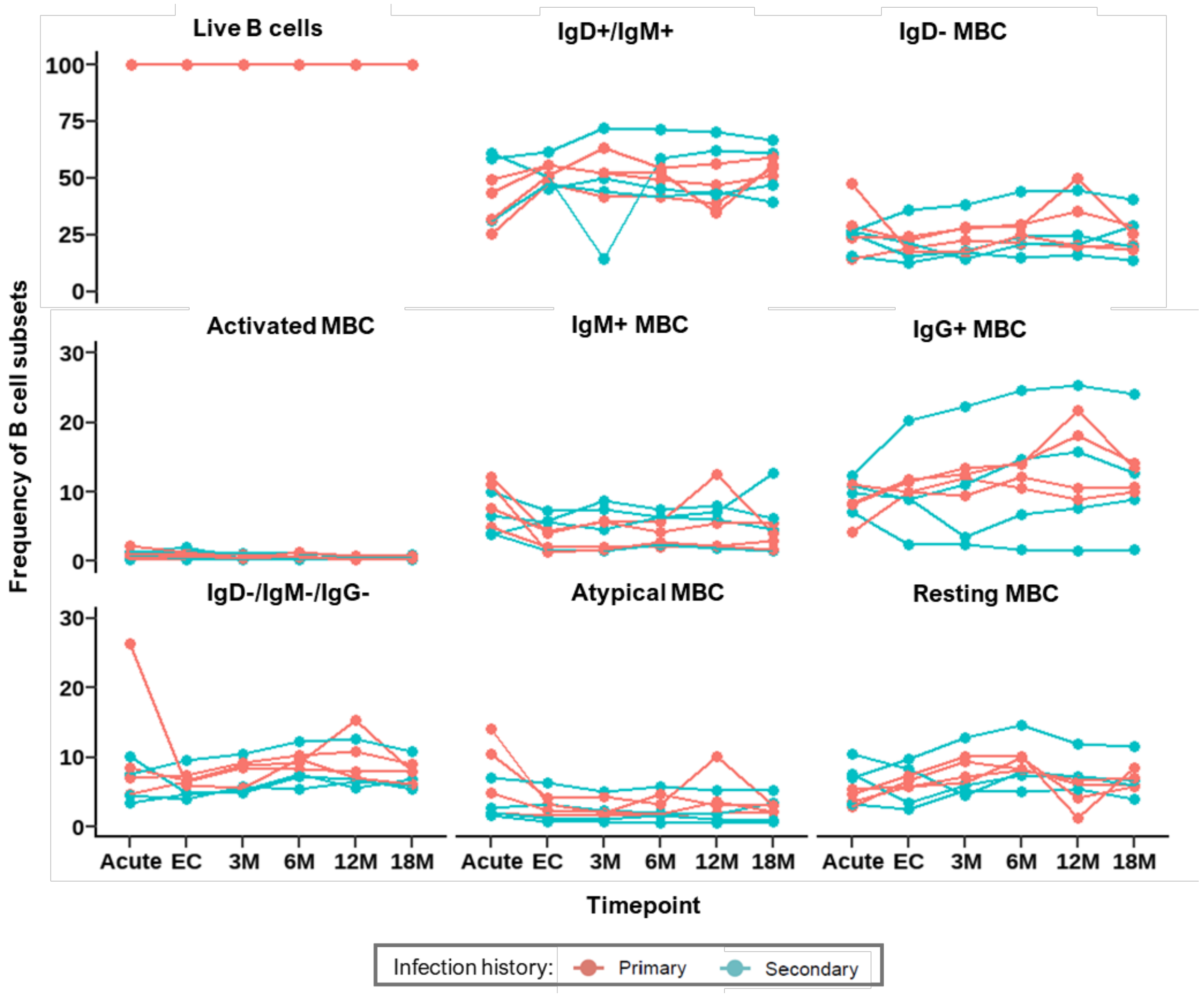


**Fig S11. <u>Total</u> B cell subset frequency as a percentage of live B cells over time.** Analytic dataset of longitudinal B cell subsets frequencies (i.e., not just the DENV-specific B cells) as a percentage of live B cells for each of the nine key B cell subsets evaluated in this study. Each solid line represents data from a single participant from either the 1° (n=4 participants; pink) or 2° (n=4 participants; blue) dengue group. Participants were sampled over time at acute, EC, 3M, 6M, 12M and 18M. B cell subsets abbreviated as follows: IgD-/IgG-/IgM- (inferred IgA), IgD- MBC (class-switched MBCs), IgG+ MBC (IgD-/IgG+), IgM+ MBC (IgD-/IgM+) and IgD+/IgM+ (naïve-like B cells).

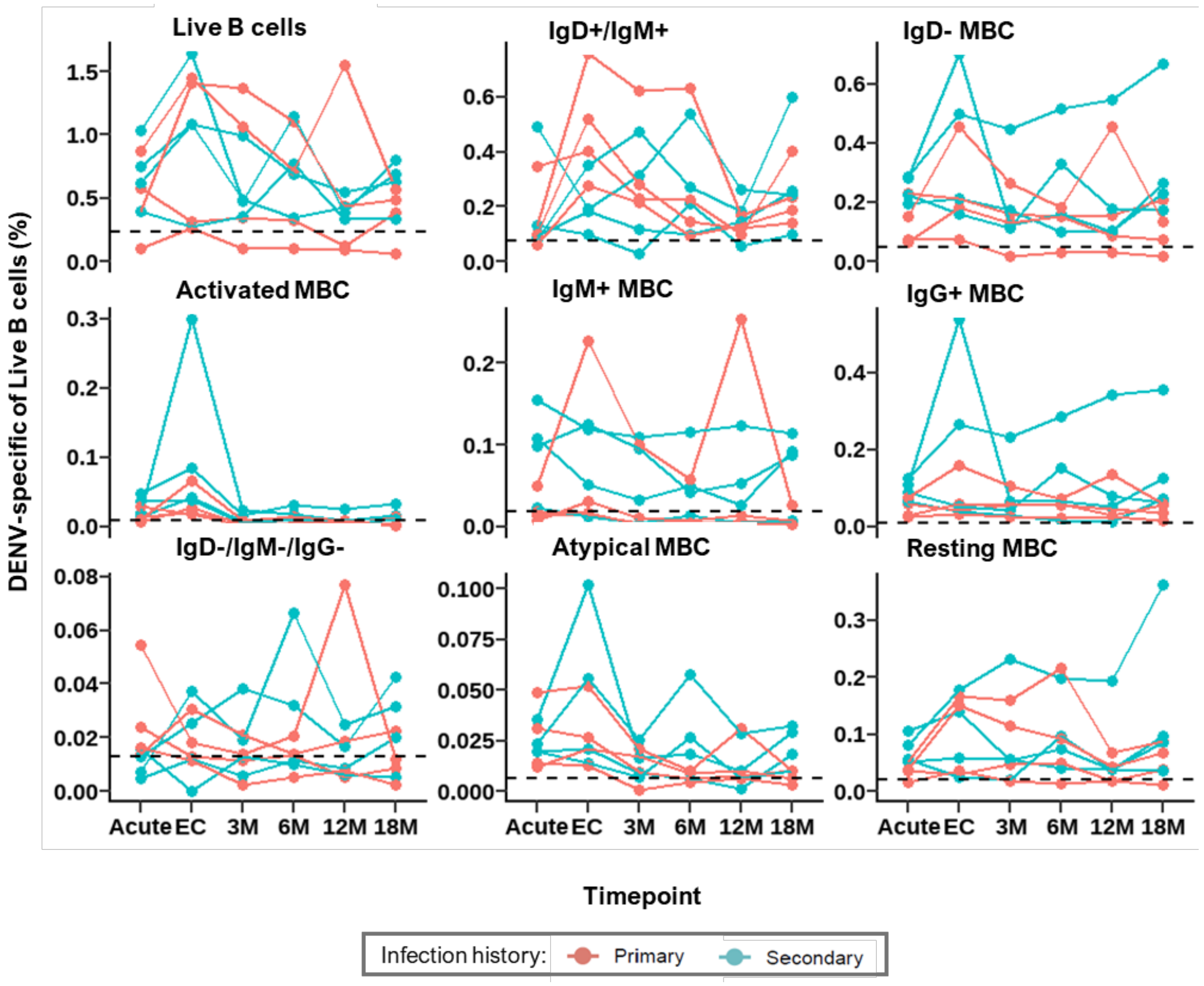


**Fig S12. Longitudinal trajectory of DENV-specific B cell frequency <u>as a percentage of live B cells</u> over time.** Analytic dataset of longitudinal DENV-specific B cell frequencies as a percentage of live B cells for each of the nine key B cell subsets evaluated in this study. A percentage of live B cells is an <u>absolute</u> metric that allows for frequency comparisons across subsets. Each solid line represents data from a single participant from either the 1° (n=4 participants; pink) or 2° (n=4 participants; blue) dengue group. Participants were sampled over time at acute, EC, 3M, 6M, 12M and 18M. DENV-specific B cell subsets abbreviated as follows: IgD-/IgG-/IgM- (inferred IgA), IgD- MBC (class-switched MBCs), IgG+ MBC (IgD-/IgG+), IgM+ MBC (IgD-/IgM+) and IgD+/IgM+ (naïve-like B cells). Dotted line indicated limit of detection based on mean PBMC samples without antigen (n=7).

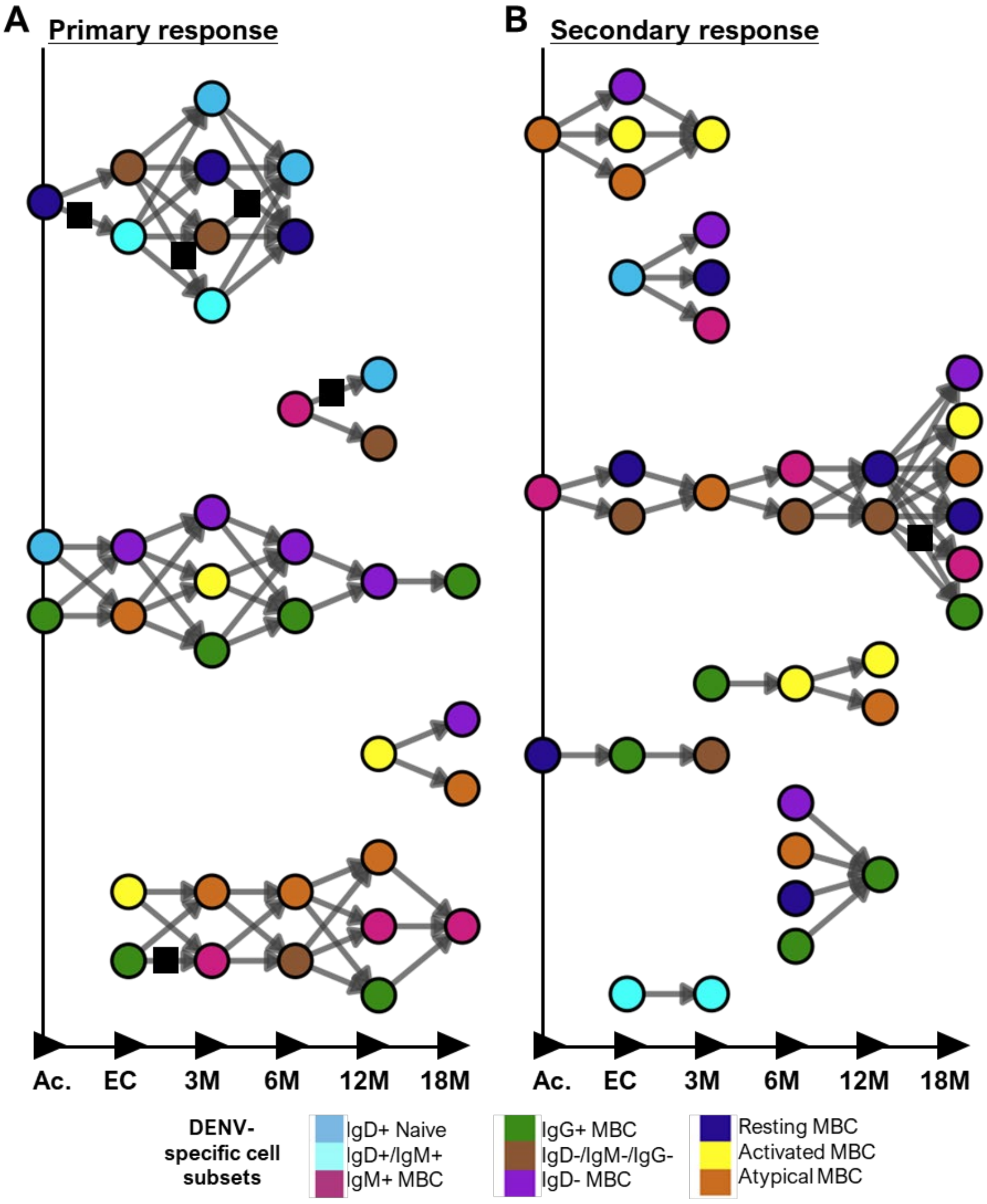


**Fig S13. Direct correlations over consecutive time bins in primary and secondary <u>DENV-specific</u> B cell frequencies <u>without exclusion</u> of biologically implausible cellular transitions.** Direct Spearman Rank correlations of DENV-specific B cell frequencies per 10,000 parental cells between consecutive timepoints (i.e., Acute-EC, EC-3M, 3M-6M, 6M-12M, and 12M-18M) at $p<0.1$ were plotted as an arrow on a directed network graph for (**A**) 1° (n=4 participants) and (**B**) 2° (n=4 participants) dengue groups. For comparison, all correlations are plotted, without applying the exclusion criteria for biologically implausible cellular transitions. Biologically implausible transitions as per Table S4 are indicated with a black square atop the arrow. Colored circles indicate DENV-specific B cell subset identity as per legend.

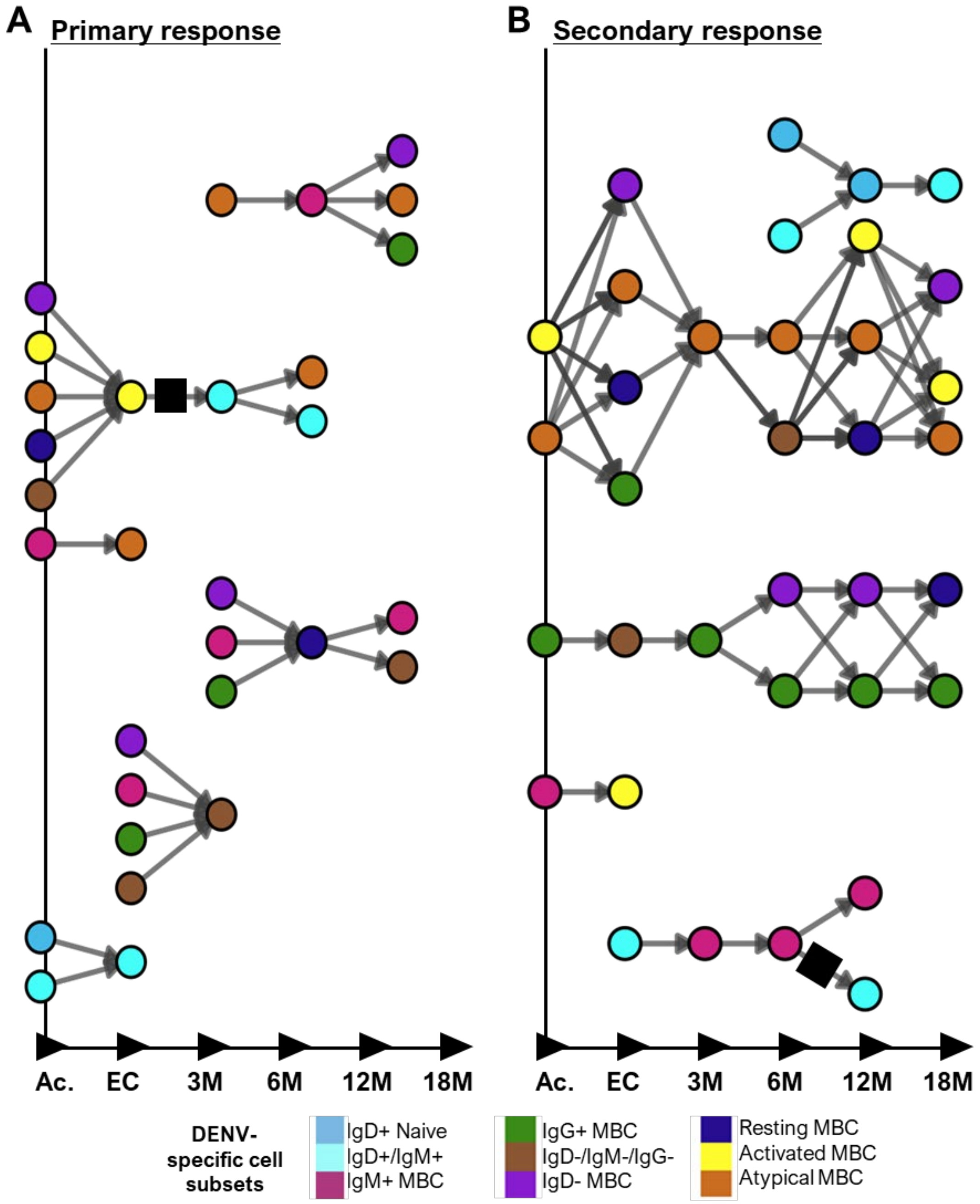


**Fig S14. Direct correlations over consecutive time bins in primary and secondary <u>total</u> B cell frequencies <u>without exclusion</u> of biologically implausible cellular transitions.** To assess whether the correlations observed in the DENV-specific B cell compartment were merely an attribute of temporal changes in frequency of total B cells (not just the DENV-specific B cells), the correlations of total B cell subset frequency were tested. Direct Spearman Rank correlations of B cell frequencies as a percentage of live B cells between consecutive timepoints (i.e., Acute-EC, EC-3M, 3M-6M, 6M-12M, and 12M-18M) at $p<0.1$ were plotted as an arrow on a directed network graph for (**A**) 1° (n=4 participants) and (**B**) 2° (n=4 participants) dengue groups. For comparison, all correlations are plotted, without applying the exclusion criteria for biologically implausible cellular transitions. Biologically implausible transitions as per Table S4 are indicated with a black square atop the arrow. Colored circles indicate B cell subset identity as per legend. Any correlations (n=11) observed in both total B cell frequencies and DENV-specific B cells were annotated on Main Figure 4.
